# Disruption of CAD oligomerization by pathogenic variants

**DOI:** 10.1101/2024.07.30.605799

**Authors:** Francisco Del Caño-Ochoa, Lobna Ramadane-Morchadi, Lluís Eixerés, María Moreno-Morcillo, Rafael Fernández-Leiro, Santiago Ramón-Maiques

## Abstract

CAD is a multi-enzymatic protein essential for initiating the de novo biosynthesis of pyrimidine nucleotides, forming large hexamers whose structure and function are not fully understood. Defects in CAD result in a severe neurometabolic disorder that is challenging to diagnose. We developed a cellular functional assay to identify defective CAD variants, and in this study, we characterized five pathogenic missense mutations in CAD’s dihydroorotase (DHO) and aspartate transcarbamylase (ATC) domains. All mutations impaired enzymatic activities, with two notably disrupting the formation of DHO dimers and ATC trimers. Combining crystal structures and AlphaFold predictions, we modeled the hexameric CAD complex, highlighting the central role of the DHO and ATC domains in its assembly. Our findings provide insight into CAD’s stability, function, and organization, revealing that correct oligomerization of CAD into a supramolecular complex is required for its function in nucleotide synthesis and that mutations affecting this assembly are potentially pathogenic.

## INTRODUCTION

Genome-wide sequencing is critical for identifying causal mutations in inherited metabolic diseases, but interpreting the impact of individual genomic alterations poses significant challenges. Many genetic missense variants have uncertain effects and require functional assays and detailed characterization of the target proteins to assess their clinical relevance and to understand the dysfunction mechanisms of the “diseased protein.”

Recent studies have identified defects in CAD, a multi-enzymatic protein initiating *de novo* pyrimidine nucleotide biosynthesis^1–3^, as the cause of an autosomal recessive neurometabolic disorder (OMIM #616457) that manifests in newborns and young children^4,5^. The disease is characterized by early-onset refractory epilepsy, developmental delay, severe developmental regression, and anemia^6^. The untreated disease is often fatal, but patients show remarkable improvement with oral supplements of uridine, which fuel pyrimidine nucleotide synthesis via a CAD-independent salvage pathway^7–16^. However, diagnosis is challenging due to symptoms overlapping with other conditions, the absence of a biomarker, and the observation of >2,400 missense CAD variants in the general population (gnomAD v4.0, ENSG00000084774). Our limited understanding of CAD complicates the prediction of whether a clinical missense variant is pathogenic or benign.

CAD is a poorly understood protein of 2.225 amino acids, formed by the fusion of four enzymatic domains, each catalyzing an initial step in *de novo* pyrimidine synthesis (Fig. 1a)^2,3,17^. This large protein further oligomerizes into hexamers of ∼1.5 MDa, nearly half the size of a ribosome^2,18^. Although the structure of the hexamer has not been described, our group determined the crystal structures of CAD’s isolated dihydroorotase (DHO) and aspartate transcarbamoylase (ATC) domains, showing that they assemble into dimers and trimers, respectively^19–22^. Combined with mutagenesis analysis, these findings support a model in which CAD trimers, linked by the ATC domains, further dimerize through the DHO domains, forming hexamers or “dimer of trimers”^23,24^. However, the architecture and functional implications of this mega-enzyme remain uncharacterized. To overcome the challenges of evaluating the pathogenicity of missense CAD variants, we developed a functional assay using a CAD-knockout (KO) human cell line that can only grow in media supplemented with uridine^25^. We transiently transfect these cells with a CAD cDNA containing the specific patient variant and monitor proliferation over a week in uridine-free conditions. This method identifies mutations that disrupt CAD activity and impede proliferation, indicating pathogenicity, while normal cell growth suggests that the variant is likely benign. This assay has been instrumental in diagnosing CAD-deficient patients worldwide^25,26^.

**Figure 1.**
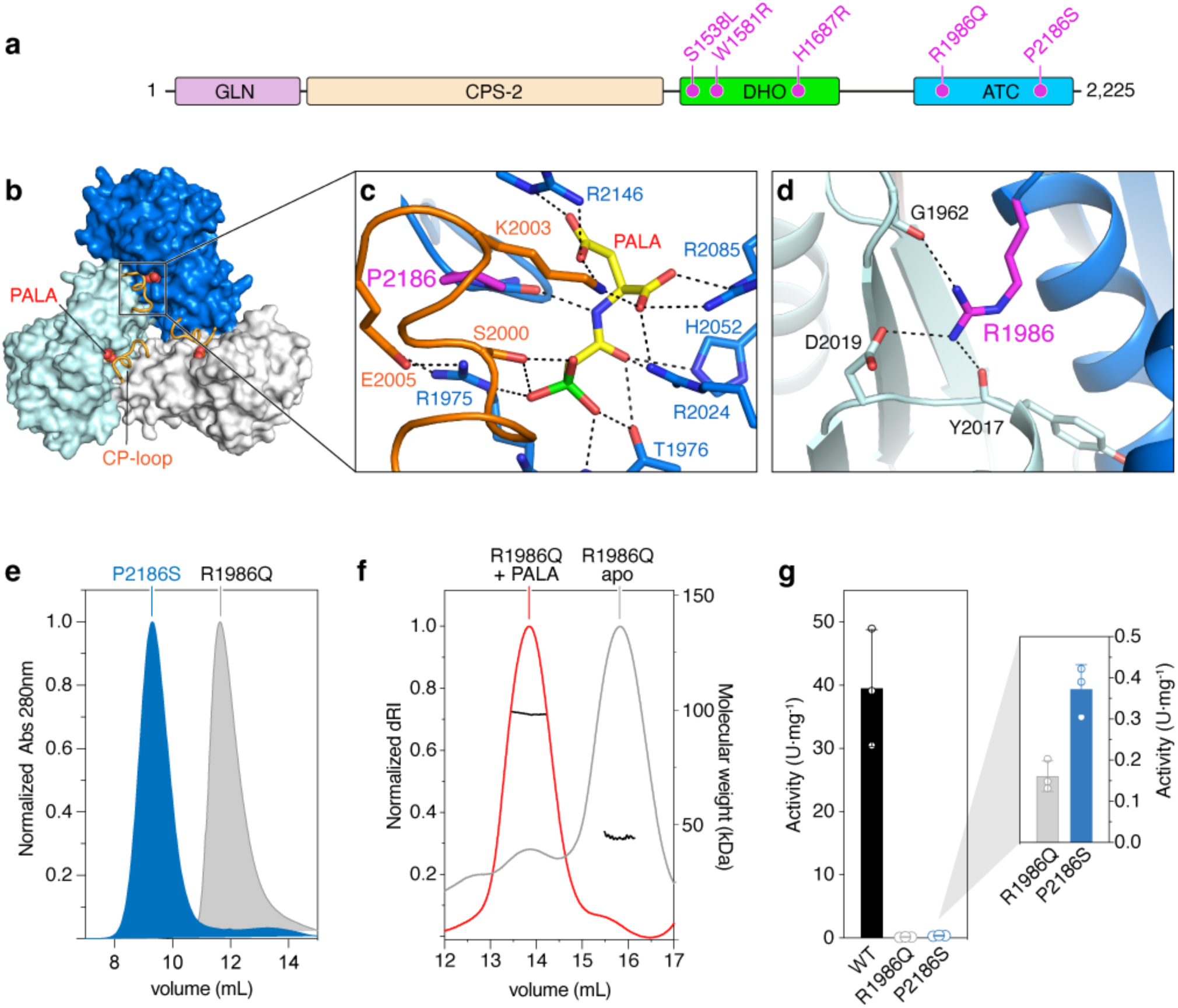
Pathogenic variants in CAD’s ATC domain. **a** Scheme of CAD protein with the four enzymatic domains represented by boxes. Pathogenic variants in the DHO and ATC domains are indicated in magenta. **b** Surface representation of CAD’s ATC homotrimer with each subunit in a different color. The CP-loop is represented in orange cartoon, and PALA as spheres. **c** Detail of PALA binding in the active site and the mutated P2186 depicted in magenta. Dashed lines represent electrostatic interactions. **d** Detail of the intersubunit interactions between the mutated residue R1986 and the neighboring subunit. **e** SEC elution of ATC mutants P2186S and R1986Q indicate the formation of trimers and monomers, respectively. **f** SEC-MALS analysis shows that PALA binding favors the assembly of R1986Q into trimers (98.4 ± 0.4 kDa), while the apo protein elutes as a monomer (39.5 ± 0.7 kDa). **g** Enzymatic activity of WT and ATC mutants shown in scattered plot with three independent measurements. Error bars indicate standard deviation. Activity units (U) are nmol of carbamoyl aspartate per min.

At the same time, identifying pathogenic mutations serves as a key to unlocking our understanding of CAD, as they point to crucial elements for the protein’s stability, function, and organization. In this study, we performed the biochemical and structural characterization of five missense mutations in CAD that were classified as pathogenic based on the cell proliferation assay and mapped within the DHO and ATC domains (Fig. 1a)^26^. Our findings reveal that although all mutations have a detrimental effect on enzymatic activity, two of them strongly impair the formation of the DHO dimers and ATC trimers. Combining AlphaFold predictions with crystal structure information, we constructed a structural model of the CAD hexamer that highlights the central role of the DHO and ATC in the architecture of the particle. This study marks the discovery of pathogenic changes that affect CAD oligomerization, underscoring the critical role of assembling into a functional supramolecular complex for the proper function of this multi-enzymatic protein in the nucleotide synthesis pathway.

## RESULTS

### Characterization of pathogenic variants in CAD’s ATC domain

The cell proliferation assay identified two pathogenic changes, R1986Q and P2186S, located within CAD’s ATC domain (Fig. 1a)^25^. This domain self-assembles into a homotrimer with three active sites at the interface between adjacent subunits (Fig. 1b)^21^. Mutation P2186S affects an invariant residue within the active site that introduces a kink in the polypeptide chain and contacts the side chains of three also invariant residues –S2000, K2003 and E2005– in the carbamoyl phosphate (CP) binding loop of the adjacent subunit (Fig. 1c and Supplementary Fig. 1). A substituting Ser at this position was predicted to distort the active site and hinder the approach of the CP-loop. On the other hand, residue R1986, located ∼20 Å away from the active site and on the opposite face of the trimer, forms an intersubunit salt bridge with D2019 (Fig. 1d). Both residues are highly conserved (Supplementary Fig. 1); thus, mutation R1986Q was anticipated to disrupt the conserved interaction and destabilize the trimer.

To characterize the molecular alterations of these variants, we produced the ATC domain bearing the point mutations R1986Q and P2186S. The recombinant proteins were produced at a similar yield and purity as the wild-type (WT) protein, suggesting that the mutations did not have a noticeable effect on protein folding and stability. SEC analysis showed that ATC-P2186S eluted at the expected position for a homotrimer (expected molecular weight 105 kDa), whereas ATC-R1986Q behaved as a monomer (expected 35 KDa) (Fig. 1e). Using SEC coupled with multiangle light scattering (MALS) analysis, we observed that adding PALA (phosphonacetyl-L-aspartate), an inhibitor that mimics the transition state of the reaction^27^, prompted the re-assembly of ATC-R1986Q into trimers, indicating that the multiple electrostatic interactions with the ligand compensated for the missing salt-bridge between subunits (Fig. 1c,f).

ATC can only be active as a trimer, and thus, the oligomerization defect of ATC-R1986Q explained the almost null enzymatic activity (Fig. 1g). Remarkably, the residual activity of this mutant was lower than that of ATC-P2186S, which, as mentioned earlier, replaces a central invariant residue within the active site.

Overall, these results indicated that R1986Q’s pathogenicity is caused by its impediment to assembling a functional ATC trimer and that this defect could be comparable to or perhaps even more harmful than other mutations, such as P2186S, with a direct impact at the active site.

### Characterization of pathogenic variants in CAD’s DHO domain

The cell proliferation assay identified three pathogenic mutations that affected conserved residues in the DHO domain: S1538L, W1581R, and H1687R (Fig. 1a and Supplementary Fig. 2)^26^. This domain forms a homodimer, with each subunit folding in an (α/β)_8_-barrel with an adjacent β-stranded region opposite the dimerization interface and a long C-terminal extension (Fig. 2a)^19,24^. The active site, formed by the loops connecting the carboxy end of the β-barrel with the surrounding α-helices, contains three Zn^2+^ cations and a flexible loop that closes over the substrate carbamoyl aspartate (Ca-Asp) and opens for dihydroorotate product release (Fig. 2a,b). A preliminary structural analysis suggested that S1538L would disrupt the formation of hydrogen bonds in the loop bearing the mutation (loop-3), which is adjacent to the catalytic flexible loop (Fig. 2b). On the other hand, W1581R replaces the Trp at the C-end of a dimerization helix with a positively charged side chain that would be in an unfavorable hydrophobic environment and cause severe steric clashes (Fig. 2c), whereas H1687R introduces a bulky side chain next to a Zn-coordinating Asp (D1686), likely causing steric clashes and distortion of the active site (Fig. 2d). The pathogenicity of the variants was correctly predicted by FoldX, with scores being two-fold higher for W1581R and H1687R compared to S1538L, whereas AlphaMissense considered S1538L as benign (Supplementary Table 1).

**Figure 2.**
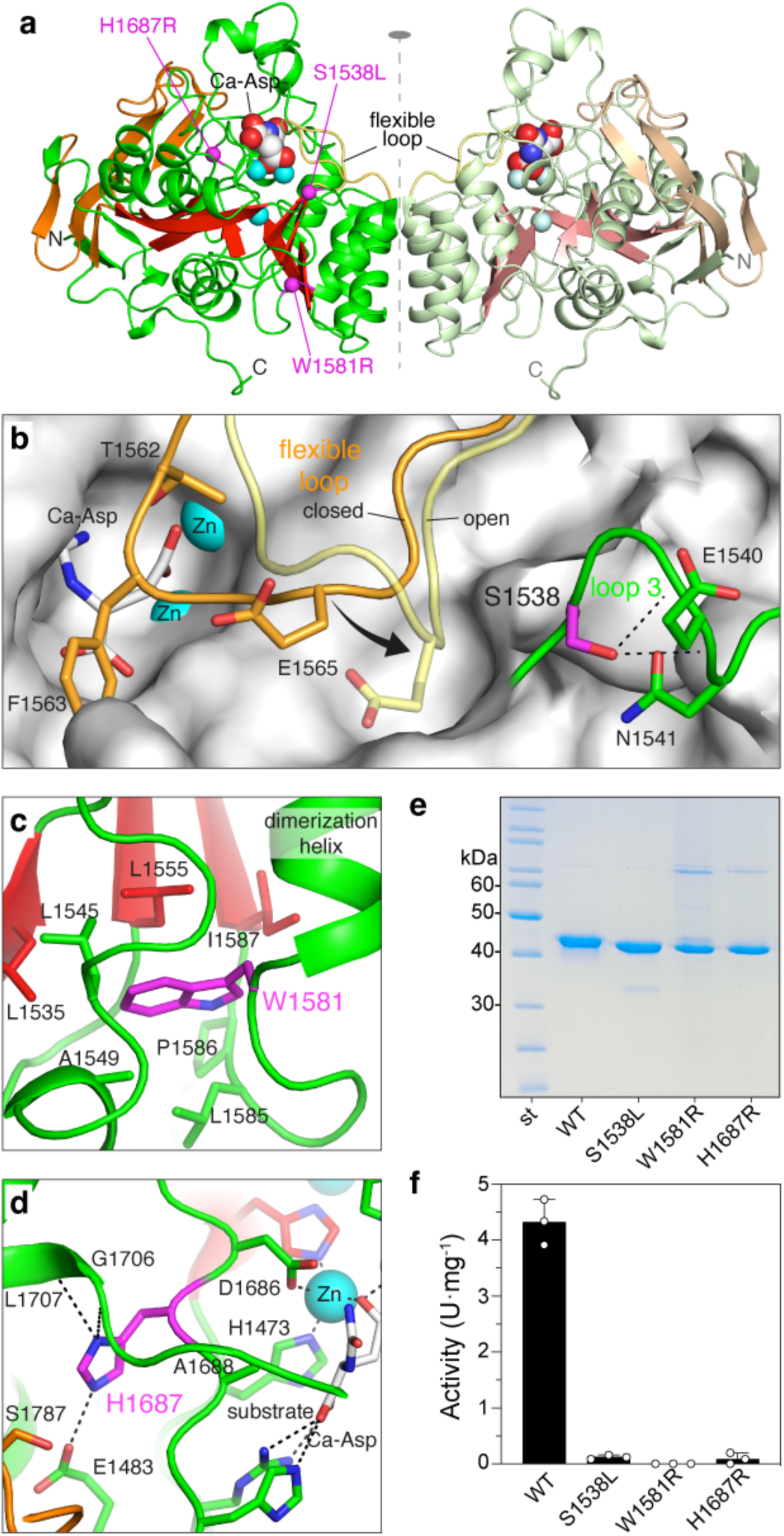
Pathogenic variants in CAD’s DHO domain. **a** Cartoon representation of the DHO dimer, with substrate Ca-Asp and Zn^2+^ cations depicted as spheres. The positions of pathogenic variants are indicated with magenta spheres. **b** Surface representation of the DHO active site with bound Ca-Asp and the flexible loop in closed (orange) and open (yellow) conformations. Loop-3 is shown in green, with the side chain of S1538 in magenta. Dashed lines indicate electrostatic interactions. **c,d** Detail views of the mutated residues (in magenta) and surrounding elements. **e** SDS-PAGE of purified recombinant DHO mutants. **f** Enzymatic activity of WT and DHO mutants shown in a scattered plot with three independent measurements. Error bars indicate standard deviation. Activity units (U) are nmol of dihydroorotate per min.

To characterize the molecular alterations of these variants, we produced the isolated DHO domains bearing the point mutations (Fig. 2e). Mutant DHO-S1538L was expressed and purified with a similar yield as the DHO-WT. In contrast, mutants DHO-W1581R and DHO-H1687R consistently exhibited ∼20-fold lower yields, indicating that these substitutions adversely affect protein folding and stability.

All three mutated DHO domains showed impaired enzymatic activity (Fig. 2f). The residual activity of DHO-S1538L (v= 0.1 U·mg^-1^) was ∼40-fold lower than the WT (v=4.2 U·mg^-1^), whereas DHO-W1581R and DHO-H1687R exhibited no activity despite testing protein concentrations 10-fold higher than the WT.

Analysis of the oligomeric state by size exclusion chromatography (SEC) showed that similar to the WT, DHO-H1687R eluted as a single peak at the position expected for a homodimer (Fig. 3a). In turn, DHO-S1538L and DHO-W1581R eluted in two peaks, corresponding to a different proportion of dimers and monomers. Mutant DHO-W1581R behaved mostly as a dimer with a 30% monomer fraction, whereas DHO-S1538L was mostly dissociated (80%) (Fig. 3a).

**Figure 3.**
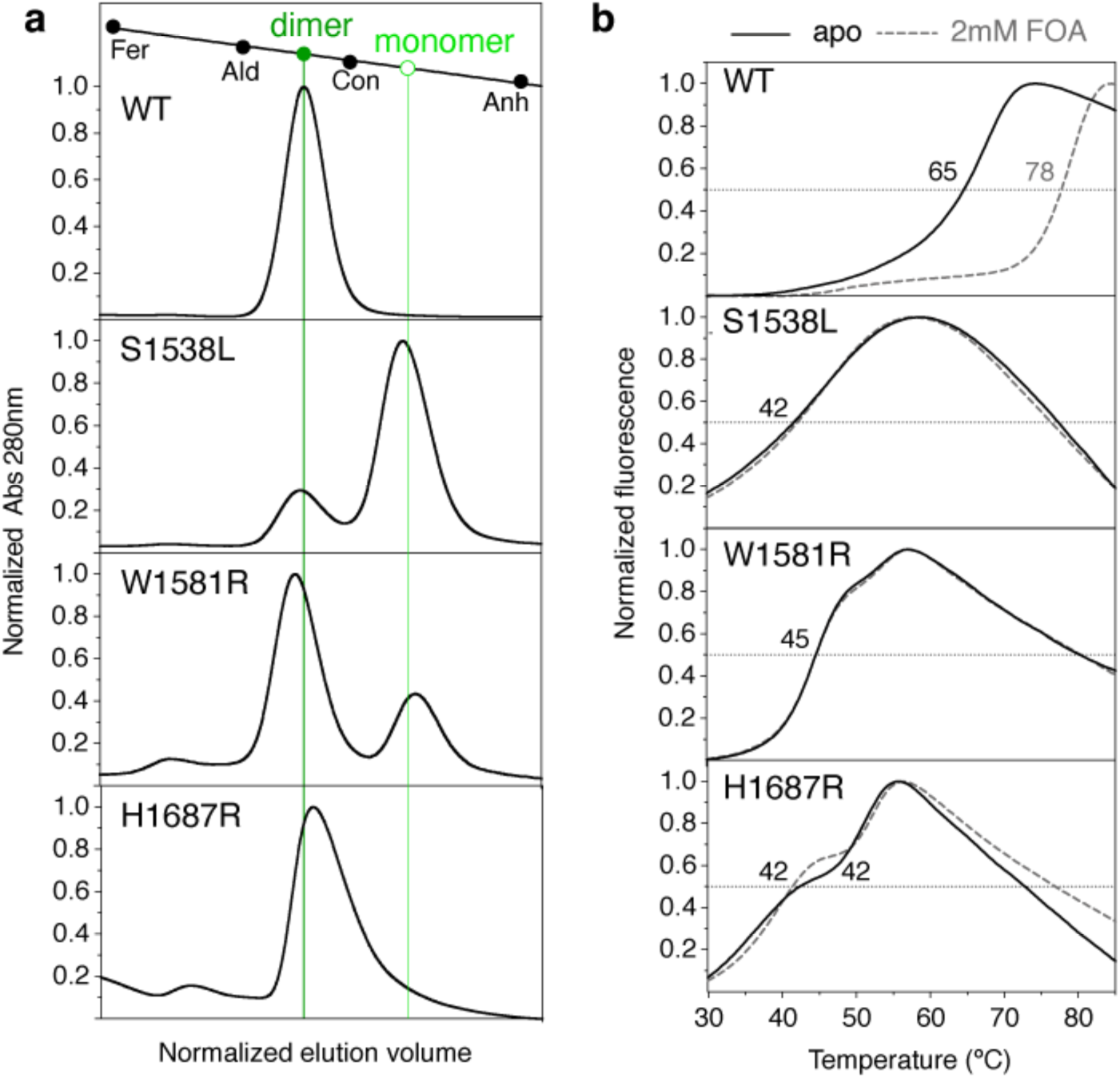
Oligomerization and stability of DHO mutants. **a** SEC analysis of purified DHO proteins. Column calibration with proteins of known molecular weight: ferritin (440 kDa), aldolase (158 kDa), conalbumin (75 kDa), and carbonic anhydrase (29 kDa). **b** Denaturing curves in the absence (apo) and presence of fluoroorotate (FOA) measured by scanning fluorimetry. Midpoint temperatures (T_m_) are indicated.

We compared the stability of the proteins by differential scanning fluorimetry (DSF) in the absence and presence of fluoroorotic acid (FOA), a competitive inhibitor that mimics the binding of dihydroorotate to the active site^19^. The DHO-WT denaturation curves showed a midpoint temperature (Tm) of 65 °C and an increment of 13 °C upon binding to FOA (Fig. 3b). In turn, the mutants had compromised stability, with Tm’s falling within the range of 42–45 °C and no increase in stability in the presence of FOA.

In summary, these results indicated that reduced stability, compromised substrate binding and impaired enzymatic activity may contribute to the pathogenicity of the mutations in the DHO domain. Notably, two mutations, S1538L and W1581R, also promoted the dissociation of the DHO dimer, suggesting that this was a novel factor to consider when evaluating the pathogenicity of CAD clinical variants.

### S1538L hinders the movement of the DHO catalytic flexible loop

To better understand the impact of the pathogenic mutations in the DHO domain, we sought to determine their crystal structures. Mutants DHO-W1581R and DHO-H1687R could not be concentrated without significant protein losses and did not crystallize in the conditions reported for the WT or after extensive screenings, supporting the destabilizing effects of these mutations. However, DHO-S1538L crystallized using the established conditions for the WT, and the structure was determined at 1.5 Å resolution (Table 1).

**Table 1.** Data collection and refinement statistics. Values for the highest resolution shell are shown in parentheses.

|  | S1538L+Ca-Asp | S1538A+Ca-Asp | S1538A apo |
| --- | --- | --- | --- |
| PDB deposition ID | 9FS1 | 9FS2 | 9FS3 |
| Data collection |  |  |  |
| Wavelength | 0.87313 | 0.97926 | 0.97926 |
| Resolution range (Å) | 73.08-1.21 (1.26-1.21) | 47.22-1.12 (1.16-1.12) | 44.43-1.18 (1.22-1.18) |
| Space group | C222<sub>1</sub> | C222<sub>1</sub> | C222<sub>1</sub> |
| Unit cell parameters<br>a, b, c (Å)<br>$\alpha$, $\beta$, $\gamma$ (°) | 82.3, 158.7, 61.9<br>90, 90, 90 | 82.1, 158.6, 61.9<br>90, 90, 90 | 82.1, 158.5, 61.8<br>90, 90, 90 |
| Total reflections | 781,132 (24,483) | 303,746 (27,639) | 671,722 (63,734) |
| Unique reflections | 119,378 (5,546) | 153,846 (13,868) | 130,558 (12,435) |
| Multiplicity | 6.4 (6.7) | 6.5 (6.1) | 5.1 (5.0) |
| Completeness (%) | 99.2 (91.5) | 98.8 (91.1) | 98.2 (95.0) |
| Mean I / sigma(I) | 20.4 (0.8) | 7.5 (0.8) | 8.0 (0.9) |
| Wilson B-factor | 13.1 | 11.0 | 11.7 |
| R-pim | 0.049 (1.048) | 0.051 (0.963) | 0.044 (0.762) |
| CC<sub>1/2</sub> | 0.992 (0.882) | 0.998 (0.410) | 0.998 (0.433) |
| Refinement |  |  |  |
| Reflections used | 119,320 (9,691) | 152,046 (13,864) | 129,893 (12,431) |
| R-factor / R-free | 0.12 / 0.15 | 0.14 / 0.17 | 0.15 / 0.17 |
| Number of atoms | 3,284 | 3,526 | 3,360 |
| macromolecules | 2,837 | 2,962 | 2,881 |
| ligands | 32 | 93 | 63 |
| solvent | 425 | 504 | 441 |
| Protein residues | 362 | 362 | 362 |
| RMS (bonds) (Å) | 0.011 | 0.014 | 0.009 |
| RMS (angles) (°) | 1.24 | 1.32 | 1.15 |
| Ramachandran favored (%) | 95.8 | 95.5 | 95.8 |
| Ramachandran allowed (%) | 3.6 | 3.7 | 3.4 |
| Ramachandran outliers (%) | 0.6 | 0.8 | 0.8 |
| Rotamer outliers (%) | 0.7 | 0.9 | 0.0 |
| Average B-factor | 19.21 | 16.65 | 16.48 |
| macromolecules | 17.12 | 14.34 | 14.62 |
| ligands | 14.85 | 23.92 | 25.08 |
| solvent | 33.42 | 29.35 | 27.92 |

DHO-S1538L only crystallized in the presence of the substrate Ca-Asp. Although predominantly monomeric in solution, the structure revealed a homodimer with subunits indistinguishable from the WT (Fig. 4a), suggesting that the higher protein concentration and favorable packing in the crystal lattice enhanced the formation of the dimer. The high-resolution electron density maps showed the Zn^2+^ cations and residues in the active site in virtually identical positions as the WT and a molecule of Ca-Asp interacting with the flexible loop in the closed conformation (Fig. 4b). The substituting Leu exhibited well-defined electron density and, in principle, did not show any alterations that could explain the impaired enzymatic activity (Fig. 4c). However, the Leu side chain would clash against E1565 if the catalytic flexible loop adopted the open conformation (Fig. 4d,e). This observation explained that the protein only crystallized in the closed state favored by the binding of Ca-Asp. We hypothesized that in the absence of Ca-Asp, the flexible loop could move away from the active site to a position different from the canonical open conformation that destabilized the subunit and affected dimerization.

**Figure 4.**
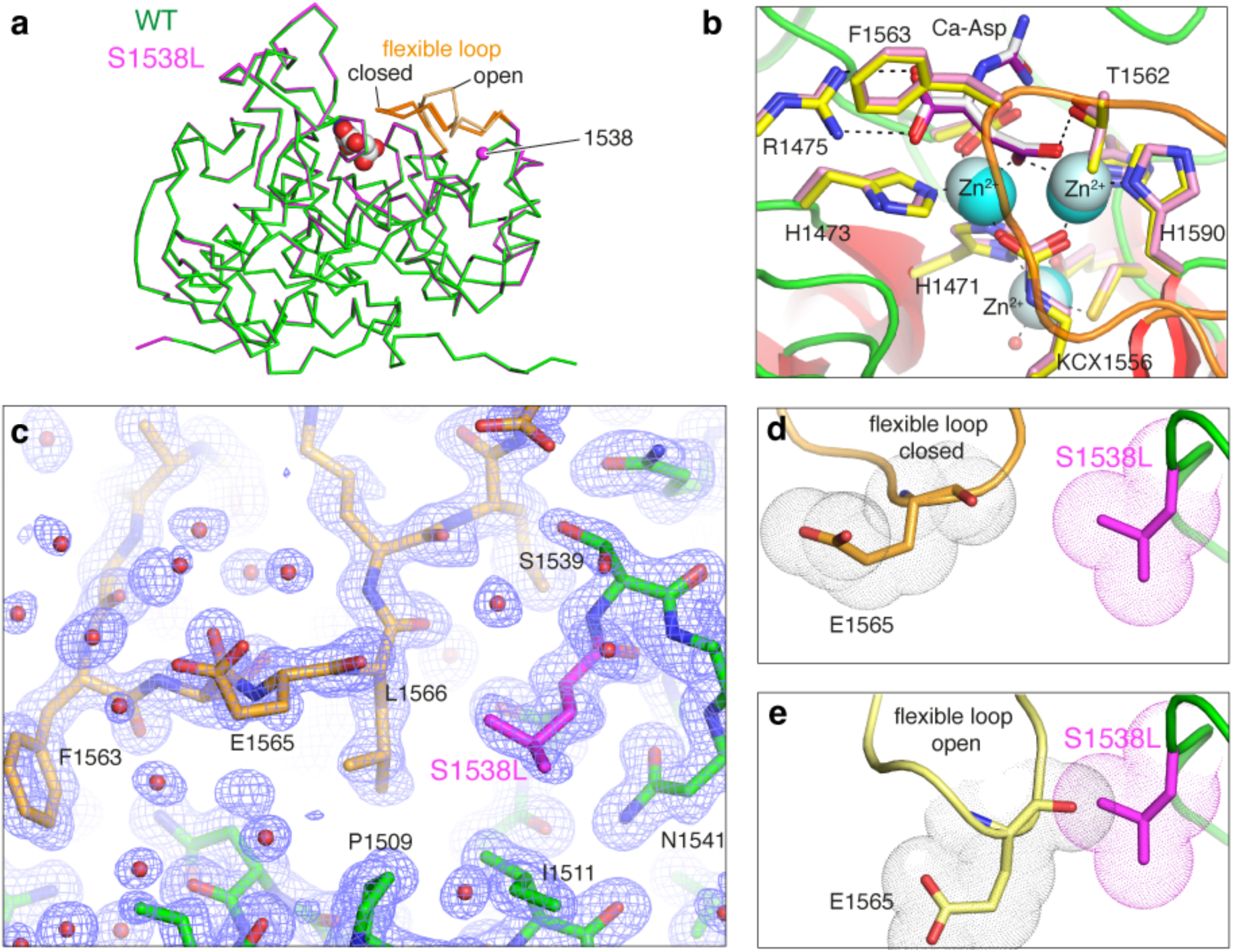
Structure of DHO S1538L mutant. **a** Ribbon superposition of DHO WT and S1538L subunits. **b** Superposition of the DHO WT (side chain carbon atoms in yellow) and mutant S1538L (pink) active sites. Dashed lines represent electrostatic interactions. **c** Detail of mutant S1538L crystal structure with the flexible loop depicted in orange and the mutated Leu in magenta. The 2F_obs_-F_calc_ electron density map is shown as a blue mesh contoured at 1.0 σ. **d** Close contact between the mutated Leu and residue E1565 in the flexible loop. Atom radii are represented as dot clouds. **e** Model of the clash between the mutated Leu and the flexible loop in the open conformation.

We also tested whether incubating mutant S1538L with 2 mM Ca-Asp and adding the substrate in the SEC column buffer favored the formation of the dimer, but the protein eluted at the expected position for a monomer (Fig. 5a). This suggested that the higher protein concentration used for crystallization and the growth of the crystal itself, which depends on the packing of a dimer in the crystal lattice, favor interdomain interactions that are otherwise too weak to maintain the S1538L dimer in solution.

**Figure 5.**
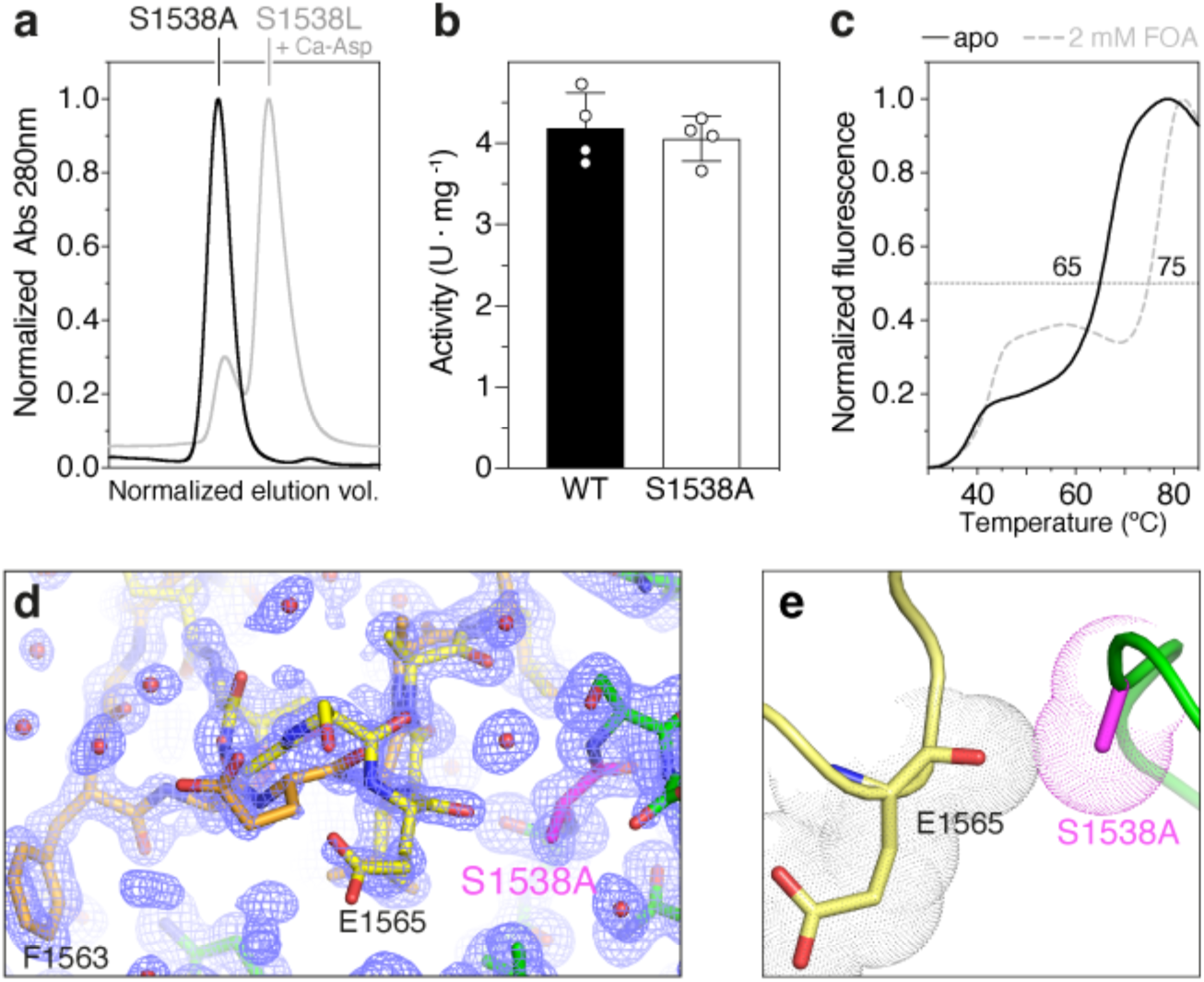
Characterization of DHO S1538A mutant. **a** SEC analysis of purified mutant S1538A (black trace) shows the formation of dimers. In the presence of 2 mM Ca-Asp, the mutant S1538L (gray trace) continues as a monomer. **b** Activity assay of mutant S1538A. **c** Thermal stability assay of mutant S1538A in the absence and presence of 2 mM FOA. **d** Detail of mutant S1538A crystal structure. The 2F_obs_-F_calc_ electron density map, shown as a blue mesh contoured at 1.0 σ, indicates a mixed population of molecules with the flexible loop in closed (orange) and open (yellow) conformations. **e** Close contact between the substituting Ala and the flexible loop, with atom radii represented as dot clouds.

To demonstrate that the inactivation of the enzyme is due to a steric clash between the mutated Leu and the loop in the open conformation, we substituted S1538 with Ala. The isolated DHO domain bearing mutation S1538A behaved as a homodimer in solution and exhibited similar enzymatic activity as the WT and comparable thermal stability (Fig. 5a–c). The structures of DHO-S1538A, free and in the presence of Ca-Asp, were determined at 1.2 Å resolution (Table 1). As for the WT, the structures showed the flexible loop in alternate open and closed conformations (Fig. 5d), and in agreement with the reversibility of the reaction, the crystal grown with Ca-Asp showed an average electron density corresponding to a mixture of substrate and product in the active site (Supplementary Fig 3). These results proved that disrupting the electrostatic interactions of the S1538 side chain within loop-3 had a minor impact on the stability and activity of the protein and that its substitution with alanine but not with leucine is compatible with the canonical open conformation of the flexible loop (Figs. 4e and 5e).

Next, we compared the movement of the flexible loop in the WT and DHO-S1538L using molecular dynamics (MD). Simulations for the protein dimers started with the flexible loop in the closed conformation and no substrate in the active site. Over the 500 ns trajectories, DHO-S1538L exhibited overall increased flexibility compared to WT, particularly in the flexible loop, the loop-3 bearing the point mutation, and the dimerization helices (Fig. 6a). Whereas in the WT, the flexible loop fluctuated between the known open and closed conformations, in DHO-S1538L, the loop moved from the closed state to distant positions that were different from the canonical open conformation. This was monitored by measuring the distance between the Cα atom of F1563 at the tip of the flexible loop relative to its position in the initially closed conformation (Fig. 6b). The increased flexibility of the mutant was also apparent by comparing the positions of the Cα atoms of residues F1563 and G1543, the residue at the tip of loop-3, along the trajectories (Fig. 6c). Principal components analysis (PCA) of the trajectories showed major conformations changes along the first eigenvector (principal component-1; PC1), corresponding to the movements of the flexible loop, loop-3 and adjacent dimerization helices (Fig. 6d). The larger variations along PC1 and PC2 indicated that mutation S1538L favored new structural states or increased the flexibility of the existing ones compared to WT.

**Figure 6.**
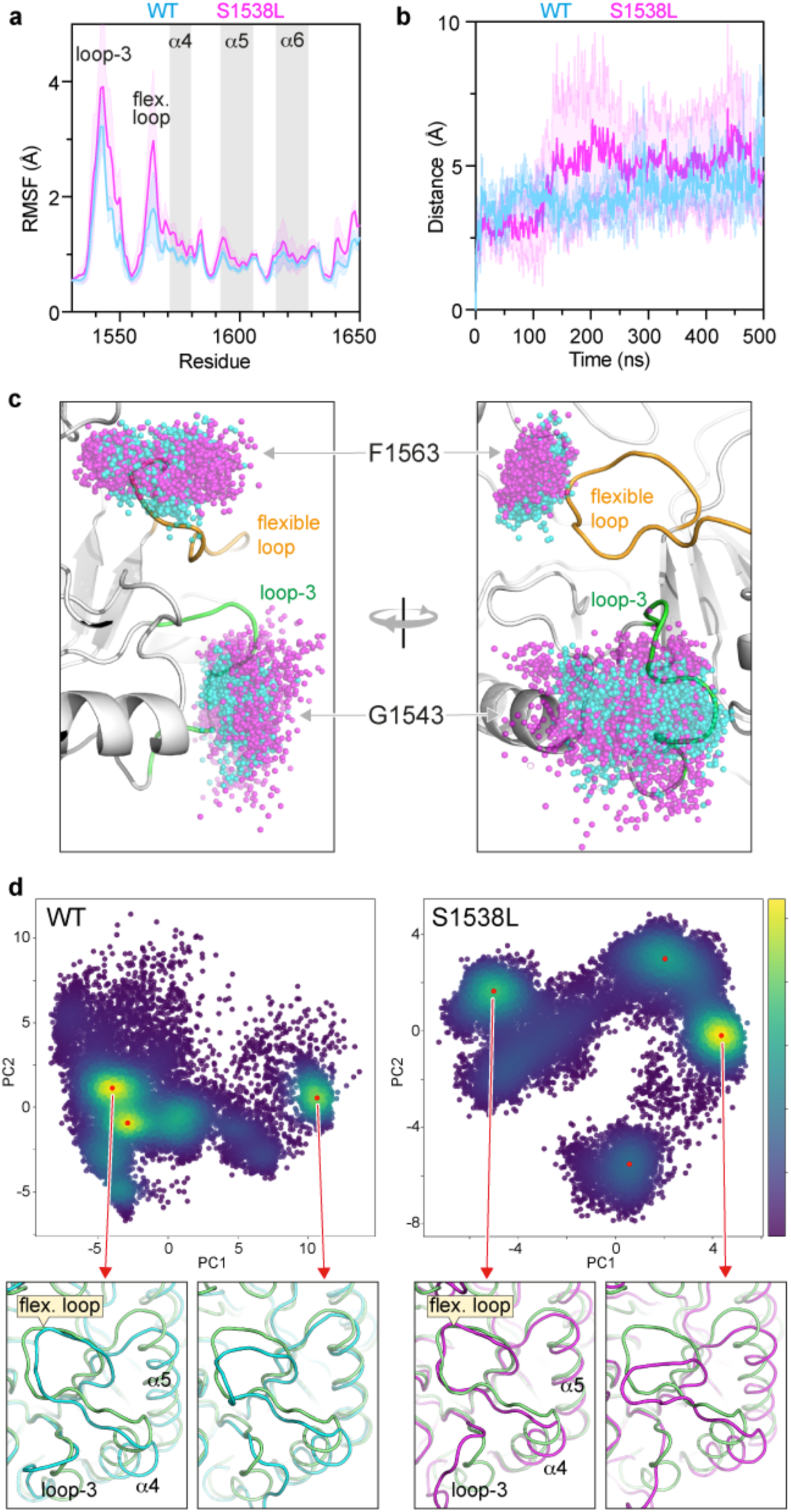
DHO molecular dynamics simulations. **a** Backbone root mean square fluctuation (RMSF) for DHO WT and mutant S1538L, showing increased flexibility in the mutant, mainly in loop-3 bearing the mutation, flexible loop and first dimerization helix (α4). Lines are mean values, and shaded areas represent the standard deviation of two independent simulations of the dimers. **b** Variation of the distance in the position of the Cα atom or residue F1563, at the tip of the flexible loop, relative to its initial closed conformation along the 500 ns trajectories. **c** Perpendicular views of the DHO model, with spheres indicating the dispersion in the positions of the Cα atoms of residues G1543 and F1563 in the WT (cyan) and mutant S1538L (magenta) during simulations. **d** Principal component analysis (PCA) of the protein models along the MD trajectories (represented as dots). Higher populated conformational states are indicated with a color gradient, with yellow being the most populated. The most populated conformations of the WT (depicted in cyan) and mutant S1538L (magenta) proteins are shown in ribbon representation and superposed with the initial closed state in the trajectory (in green).

Overall, these findings supported that mutation S1538L inactivates the enzyme by altering the movement of the catalytic flexible loop and also increases the flexibility in this loop, in loop-3 bearing the mutation, and in the adjacent helices, which may impair DHO dimerization.

### DHO dimerization is required for CAD activity

The effect of the mutations affecting DHO dimerization was at least twofold, as they also impacted the overall stability of the domain (W1581R and S1538L) and the movement of the flexible loop (S1538L). To investigate to what extent a defect in DHO dimerization contributed to the pathogenicity of the variants, we turned to another mutation, M1601E, which we previously designed to identify the DHO dimerization interface (Fig. 7a)^19^. Our prior study showed that DHO-M1601E behaved as a monomer in solution and had similar thermal stability as the WT. Interestingly, DHO-M1601E retained 50 % of the WT activity^19^, suggesting that dimerization could somehow affect the movement of the catalytic flexible loop but was not essential for the DHO enzymatic activity. However, the impact of mutation M1601E within the full-length CAD protein had not been tested.

**Figure 7.**
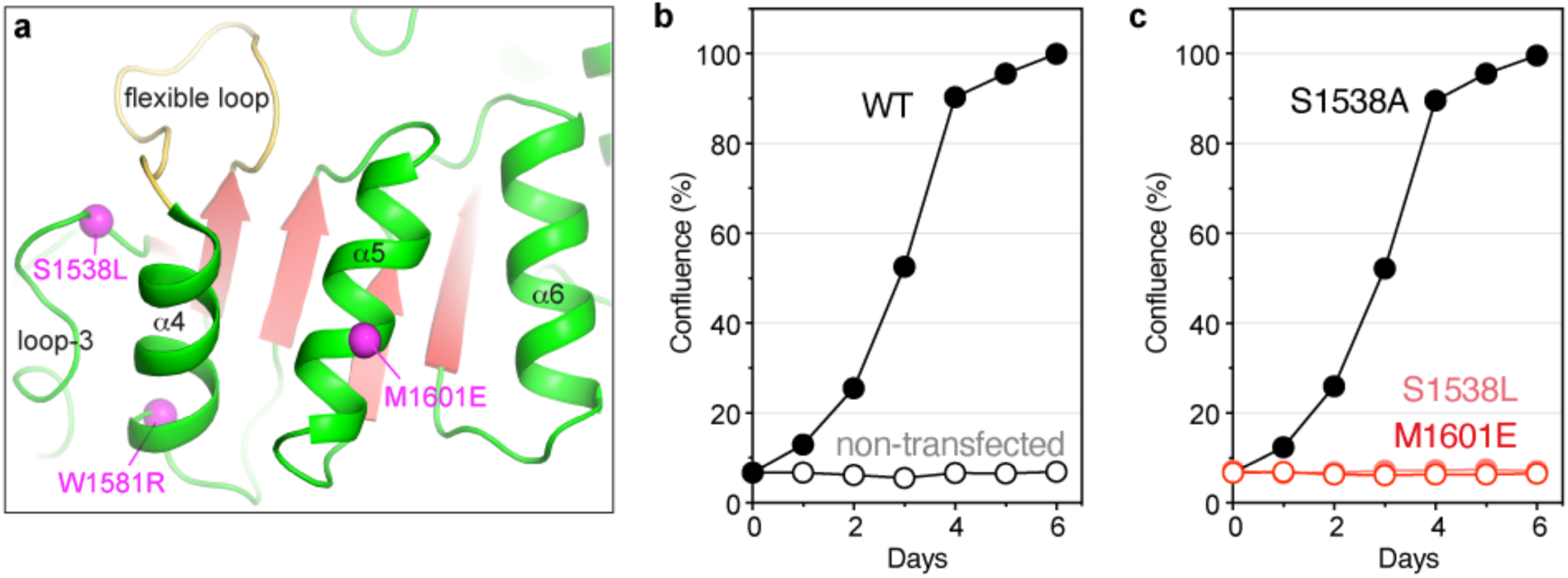
CAD is inactivated by impaired DHO oligomerization. **a** Cartoon representation of the DHO dimerization interface viewed from the other subunit in the dimer. The Cα atoms of the mutated residues are shown as magenta spheres. M1601E is located in the middle of the central dimerization helix and impedes the oligomerization by charge repulsion and steric clashes with the same residue in the other subunit. **b,c** Growth complementation assay of CAD-knockout cells grown without uridine and transfected with GFP-CAD WT (**b**) or bearing point mutations (**c**). Each point represents the mean and standard deviation of two independent experiments, counted in duplicates.

We introduced mutation M1601E in the full-length CAD construct and tested its functionality in the CAD-KO cell proliferation assay. Transfection of CAD-M1601E failed to rescue the growth of CAD-KO cells under uridine-free conditions, as seen for the pathogenic variants, including S1538L (Fig. 7b,c). This result indicated that mutation M1601E impairs CAD activity and, thus, the *de novo* synthesis of pyrimidine nucleotides. In turn, CAD-S1538A effectively recovered the growth phenotype, confirming the neutral effect of this mutation in both the isolated DHO domain and the full-length protein (Fig. 7c).

In summary, our finding with mutation M1601E indicated that DHO dimerization is essential for CAD function. Therefore, we also concluded that the deleterious effect of S1538L and W1581R on the dimerization of the DHO domain further contributes to the pathogenicity of these variants.

### Predicted structure of CAD hexamers

The low stability of human CAD protein has thus far impeded its structural characterization through various methods, including cryo-electron microscopy. Thus, we turned to the AlphaFold (AF) artificial intelligence prediction program to advance our understanding of the protein’s architecture^28^. The available CAD model in the AF database (https://alphafold.ebi.ac.uk/entry/P27708) correctly predicts the structures of the ATC and DHO domains and generates a reliable GLN/CPS-2 heterodimer, closely resembling *E. coli* CPS and human CPS-1 structures^29,30^ (Supplementary Fig. 4a). The high value for the pLDDT confidence measure (>90) indicated high accuracy in the modeled domains but not for the linkers (pLDDT <50), particularly for the 96 aa connecting DHO and ATC. However, the model fails to provide a reliable orientation of the ATC domain, as indicated by the predicted aligned error (PAE) calculations, which we attributed to the low structural complexity of the linker to the rest of the protein (Supplementary Fig. 4a).

Nonetheless, the models obtained by running AF-2 locally or using the AF-3 webserver consistently predicted the interaction between ATC and the L2 and L3 domains of CPS-2 (Fig. 8a and Supplementary Fig. 4b,c). The scores for the template modeling (pTM >0.83) and the interface predicted template modeling (ipTM = 0.76) indicated a low error in the overall predicted fold and in the relative positioning of the domains. The result was consistent whether using as input the full-length protein or the isolated enzymatic domains. In this model, the orientation of the ATC domain was compatible with its oligomerization as a homotrimer, as observed by X-ray crystallography (Fig. 1b). By superimposing three copies of the predicted CAD subunits with the crystal structure of the ATC trimer, we constructed a model of a CAD trimer that showed no steric clashes among domains (Fig. 8b). We obtained a similar model among the top five solutions of AF-2 upon modeling a CAD trimer. Within this trimer, the three DHO domains were oriented on the same side of the particle (Fig. 8c), which was compatible with the formation of the DHO homodimers observed in the crystal structure (Fig. 2a). Based on the symmetry of the DHO dimers, we positioned a second CAD trimer, obtaining a hexameric particle (a dimer of trimers) (Fig. 8d and Supplementary movie). The resulting model, obtained simply by applying the known symmetries of the ATC and DHO domains to the AF subunit solution, shows no clashes, and only small adjustments in two DHO subunits would be needed to match exactly the interactions observed in the crystal structure. It seems to us that it would be very unlikely to generate a hexameric particle without collisions in this way if the domains had a different orientation, which is an indication to us that the model must be correct.

**Figure 8.**
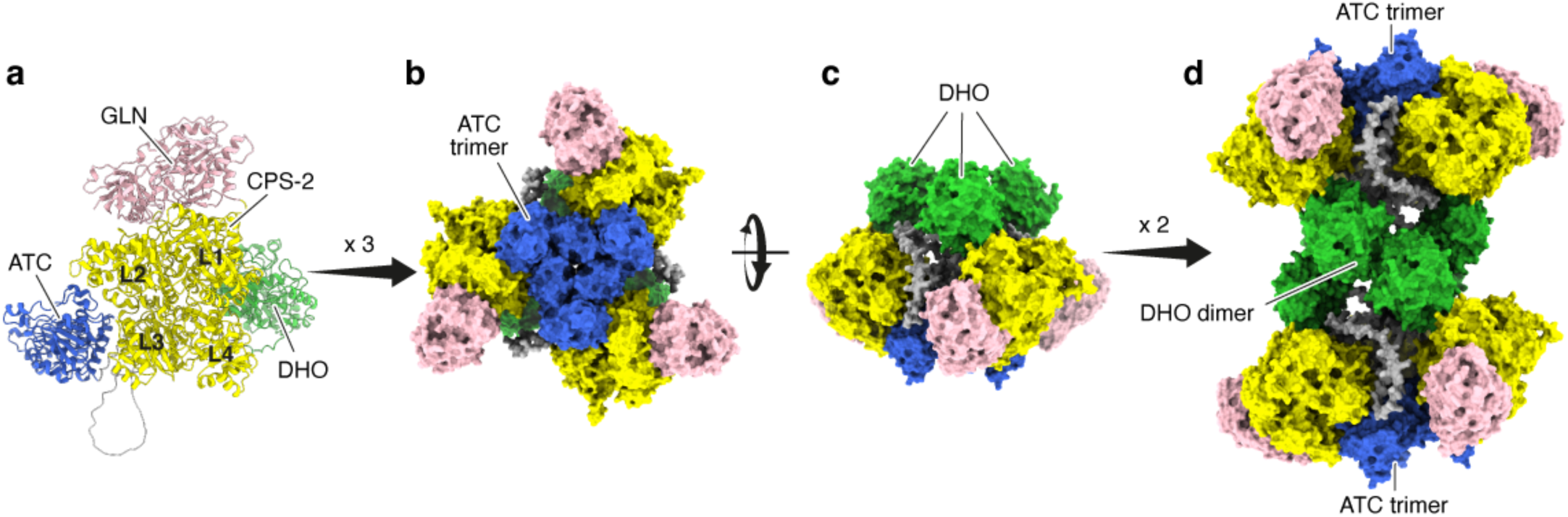
Model of CAD hexamer. **a** Cartoon representation of the model generated by AlphaFold for the CAD subunit, with each enzymatic domain depicted in a different color. **b,c** Perpendicular views of the CAD trimer obtained by superimposing three copies of the AlphaFold model on the crystal structure of the ATC trimer. **d** Model of the CAD hexamer generated by the superposition of two CAD trimers on the crystal structure of the DHO dimers.

## DISCUSSION

The assembly of CAD into a multi-functional hexameric particle has remained an unsolved puzzle with uncertain functional implications. This complex may support concatenated enzymatic reactions and regulatory mechanisms, including subunit cooperativity^31^, intermediary metabolite channeling^32–34^, allosteric control^3^, and modulation by phosphorylation^35–38^. In addition, it may be required for interactions with other proteins in the pyrimidine biosynthetic pathway, forming a metabolon known as the “pyrimidinosome”^39^. Thus, we hypothesized that mutations affecting CAD assembly might be deleterious. However, to date, the variants studied in CAD-deficient patients have primarily affected protein stability or active site elements, not oligomerization^26^. The novelty of this study is the characterization of pathogenic variants that impair the oligomerization of the DHO or ATC domains, demonstrating that the assembly of CAD into a mega-enzyme is essential for nucleotide synthesis.

The significance of ATC oligomerization in assembling an active CAD hexamer *in vitro* was first described by D. Davidson’s group^40,41^. Based on the structure of *E. coli* ATC catalytic trimer and sequence conservation, they demonstrated that disrupting the ionic pair D2009–R2187 in the ATC domain caused hamster CAD to dissociate into monomers, severely reducing enzymatic activity. They also found that adding the ATC inhibitor PALA or the substrate CP restored the CAD hexamers. We demonstrated a similar defect in the oligomerization of the ATC domain of human CAD as the pathogenic mechanism of mutation R1986Q. This mutation, found in homozygosis in a patient and one deceased sibling^25^, disrupts another conserved intersubunit ion pair (R1986-D2019), promoting the dissociation of the ATC trimer (Fig. 1e). Since the ATC active sites are formed between subunits, the monomeric domain is inactive (Fig. 1g), with the residual activity likely due to partial reassembly upon binding CP, similar to Davidson’s findings in hamster CAD. We also observed that the binding of PALA compensates for the loss of the salt bridge and restores oligomerization (Fig. 1f), indicating that the dissociation is reversible and that mutation R1986Q is not causing protein misfolding. A similar pathogenic effect can be postulated for missense mutations in D2019 that would also impair the ionic pair. According to the gnomAD database, CAD variants D2019A and D2019N appear at very low allele frequency in the population (10^-6^–10^-7^) and not in homozygosis, similar to R1986Q, which would agree with a potentially damaging effect, although they have not been found in CAD-deficient patients so far^26^.

The difficulty of producing recombinant human CAD prevented us from studying the effect of R1986Q on the oligomerization of the full-length protein. However, we previously showed that CAD-R1986Q fails to rescue the growth of CAD-KO cells without uridine^25^, leading us to conclude that the defect in ATC oligomerization inactivates CAD and impedes the synthesis of pyrimidine nucleotides. Moreover, comparing the residual activity of ATC-R1986Q and ATC-P2186S suggests that a defect in ATC oligomerization can be at least as detrimental as replacing an invariant residue within the active site (Fig. 1g).

The importance of the DHO domain for CAD assembly was less understood. The discovery that the isolated human DHO domain forms a homodimer was insightful but not entirely unexpected^19,20,42^, given that bacterial homologs form various dimeric structures^19^. We also found that the human DHO dimer was virtually identical to the pseudo-DHO domain in the CAD-like protein from fungi^24^. In fungi, GLN, CPS-2 and ATC are fused into a CAD-like polypeptide with an inactive DHO domain, while an independent monofunctional protein carries out this enzymatic activity^43^. The striking structural similarity between human and fungal DHO homodimers, along with protein-engineering assays, suggested that regardless of the enzymatic activity, DHO fulfills a conserved role in assembling CAD into a “dimer of trimers” (a hexamer)^24^. Now, the characterization of the pathogenic variants confirms that DHO dimerization is required for CAD function.

Two DHO variants, H1687R and W1581R, were found in compound heterozygosity in the first Spanish patients diagnosed with CAD deficiency^26^. These mutations were identified through post-mortem exome sequencing in one subject, and her sibling, who presented the same variants, showed favorable progression with uridine treatment (Dr. P Sanchez-Pintos, personal communication). Another DHO variant, S1538L, was identified in compound heterozygosity with an aberrant splicing variant in a 1-year-old patient who manifested symptoms from birth and improved with uridine treatment^26^ (Dr. J Kenny, personal communication). H1687R and W1581R affect the folding and stability of the isolated DHO domain, evidenced by reduced production yields, loss during concentration, decreased thermal stability, and resistance to crystallization. This instability affects the active site since both mutants lack enzymatic activity (Fig. 2f), and W1581R also causes partial dimer dissociation (Fig. 3a), likely due to its position at one end of the dimerization helix α4 (Fig. 2c and 7a). S1538L is less destabilizing than the previous mutations, as DHO-S1538L was produced and crystallized as WT, but it inactivates the enzyme and favors dimer dissociation (Figs. 2f and 3a). The DHO-S1538L structure suggested a clash with the catalytic flexible loop if this adopted an open conformation (Fig. 4c–e). In contrast, mutation S1538A did not alter the stability, activity, or dimerization of the DHO domain and did not affect the movement of the flexible loop (Fig. 5). Thus, we conclude that the altered movement of the flexible loop underlies the damaging mechanism of mutation S1538L.

The flexible loop acts as a lid over the active site in DHOs from *E. coli* and humans, playing a crucial role in the reaction cycle^44–46^. It closes to bind and orient Ca-Asp, increases its electrophilicity, excludes water molecules, stabilizes the transition state, and then opens to release dihydroorotate. Only two distinct open and closed conformation states were observed in various human DHO structures unaffected by crystal contacts^44^. Mutation S1538L blocks the canonical open conformation, forcing the catalytic loop into more distant and flexible conformations in the absence of Ca-Asp, as shown by MD simulations (Fig. 6). This alteration occurs with greater flexibility of loop-3, bearing the mutation, and the adjacent dimerization helices, possibly explaining the dimerization problems. Interestingly, we previously showed that replacing the flexible loop of human DHO with that from *E. coli* resulted in an inactive chimeric protein with impaired dimerization^44^. These findings suggest that alterations in the movement of the flexible loop can affect nearby dimerization helices, hindering DHO oligomerization.

The dual impact of S1538L on enzymatic activity and dimerization left unclear whether impaired DHO oligomerization contributes to the variant’s pathogenicity. To explore this, we examined a non-clinical mutation, M1601E, initially designed to confirm that helices α4, α5 and α6 form the DHO intersubunit surface (Figs. 2a and 7a)^19^. M1601 occupies a central position in helix α5, and the large, negatively charged side chain of the substituting Glu would clash with the same residue in the adjoining subunit, preventing dimerization. Previously, we showed that monomeric DHO-M1601E had similar stability to WT and 50% reduced activity^19^, which we could now attribute to the increased flexibility of the unrestrained dimerization helices altering the movement of the flexible loop. These results strongly suggest that, although dimerization is not essential for the activity of the isolated DHO domain, it enhances the correct movement of the flexible loop.

Besides the important link between dimerization and activity in the isolated DHO domain, mutation M1601E also helps understand the defect of DHO dimerization in the full-length protein. The difficulty in producing human CAD prevented us from studying the effect of mutation M1601E in oligomerization. Thus, we used the CAD-KO proliferation assay and proved that M1601E impedes the correct function of the full-length CAD protein (Fig. 7b,c). The 50% reduction in DHO activity caused by M1601E does not explain the damaging effect of this variant. In CAD, the rate-limiting reaction is catalyzed by the CPS-2 domain^3^, which is estimated to be ∼30-fold slower than the reaction catalyzed by the DHO domain^40^. Therefore, a twofold decrease in DHO activity is not expected to affect the overall CAD reaction, and thus, the effect of mutation M1601E can be attributed solely to impaired dimerization. We conclude that DHO dimerization is necessary to assemble a functional CAD complex, and we can also anticipate that any mutation that directly alters the dimerization interface or adjacent elements, hindering DHO oligomerization, will be pathogenic.

CAD is a complex protein to handle, and various groups (including us) have made considerable efforts to resolve its architecture by X-ray crystallography and cryo-EM, without success. To our surprise, and in a mixture of joy and dejection at the superiority of artificial intelligence, AlphaFold helped us generate a CAD structure that is likely correct, or at least very close to the real solution. Although the model requires experimental verification, it currently explains the damaging effects of the variants under study. In this model, two layers of ATC trimers surrounded by GLN/CPS-2 heterodimers are connected by three DHO dimers occupying the center of the particle (Fig. 8). The GLN domains protrude from the particle, and the CPS-2 domains only contacts the ATC and DHO domains within their own subunits. The ATC and DHO active sites face outwards, contrary to our previous hypothesis of an internal reaction chamber^24^. The exit of the CPS tunnel, which in other homologs connects the different active sites and releases the unstable ATC substrate CP^29,30^, is conveniently located near the ATC active site. The gaps between the CPS-2 domains are partially filled by the long linkers connecting the DHO and ATC domains. Phosphorylation of residue S1859 in this loop has been proposed to favor CAD oligomerization^37^, although the mechanism for this enhanced interaction remains unclear from the model. What this model certainly shows is that oligomerization of the ATC and DHO domains is essential for the formation of the CAD hexamer. In particular, the DHO domain plays (literally) a central function, which explains the accumulation of pathogenic mutations and the structural conservation in fungal CAD-like proteins despite the lack of enzymatic activity. However, the model does not explain why the hexamer is necessary for the correct activity of CAD in the cell beyond the formation of the active sites between ATC subunits. Nevertheless, our current study leaves no doubt that CAD is required to form this complex, as demonstrated by the characterization of the oligomerization defects of the pathogenic variants R1986Q and S1538L and the mutation M1601E.

Given the need to assemble a complex CAD particle, we ask whether a variant with impaired oligomerization could interfere with the function of the normal protein and thus exert a dominant negative effect. For example, the S1538L variant with impaired DHO dimerization could oligomerize with the WT protein via the ATC domain, “poisoning” the assembly of functional CAD particles (Fig. 9a,b). Similarly, mutation R1986Q impairs ATC trimerization, but the variant could still oligomerize with the WT through the DHO domain. However, the parents carrying one allele of the oligomerization-impaired CAD variants did not manifest the disease, indicating the recessive nature of these changes. This lack of a dominant effect could be due to reduced stability (observed for mutant DHOs) and degradation of pathogenic variants. Alternatively, it is plausible that only subunits with intact DHO and ATC domains are incorporated into the complex, excluding imperfect variants that cannot bind tightly. Another unexplored possibility is that mutations may weaken but not completely prevent oligomerization with WT protein (Fig. 9c). In our assays, only the mutant protein is present, as in patients with variants R1986Q (homozygosis) or S1538L (haploinsufficiency). However, in heterozygosity, mutant variants may still oligomerize with the WT protein, forming imperfect and perhaps partially functional complexes. For example, while ATC-R1986Q cannot form a homotrimer, it might form a heterotrimer with WT because the mutant subunit retains residue D2019 to form a salt bridge with residue R1986 in the WT protein (Fig. 9c). Similarly, a DHO domain with the S1538L mutation and “shaking” intersubunit helices might not dimerize with another mutant copy, but could potentially do so with a WT subunit with a solid intersubunit surface. For either of these scenarios or others we may not have considered, mutations S1538L and R1986Q are recessive. However, this does not exclude the possibility of other oligomerization defective variants being dominant. Indeed, we predict that the non-clinical mutation M1601E would be dominant as it does not diminish the stability of the DHO domain^19^ and, very likely, prevents the formation of a heterodimer with WT. Given the severity of the disease and its early manifestation, these potentially dominant mutations are expected to appear de novo and most likely in heterozygosity with an unaffected CAD variant, complicating the accurate diagnosis of patients.

**Figure 9.**
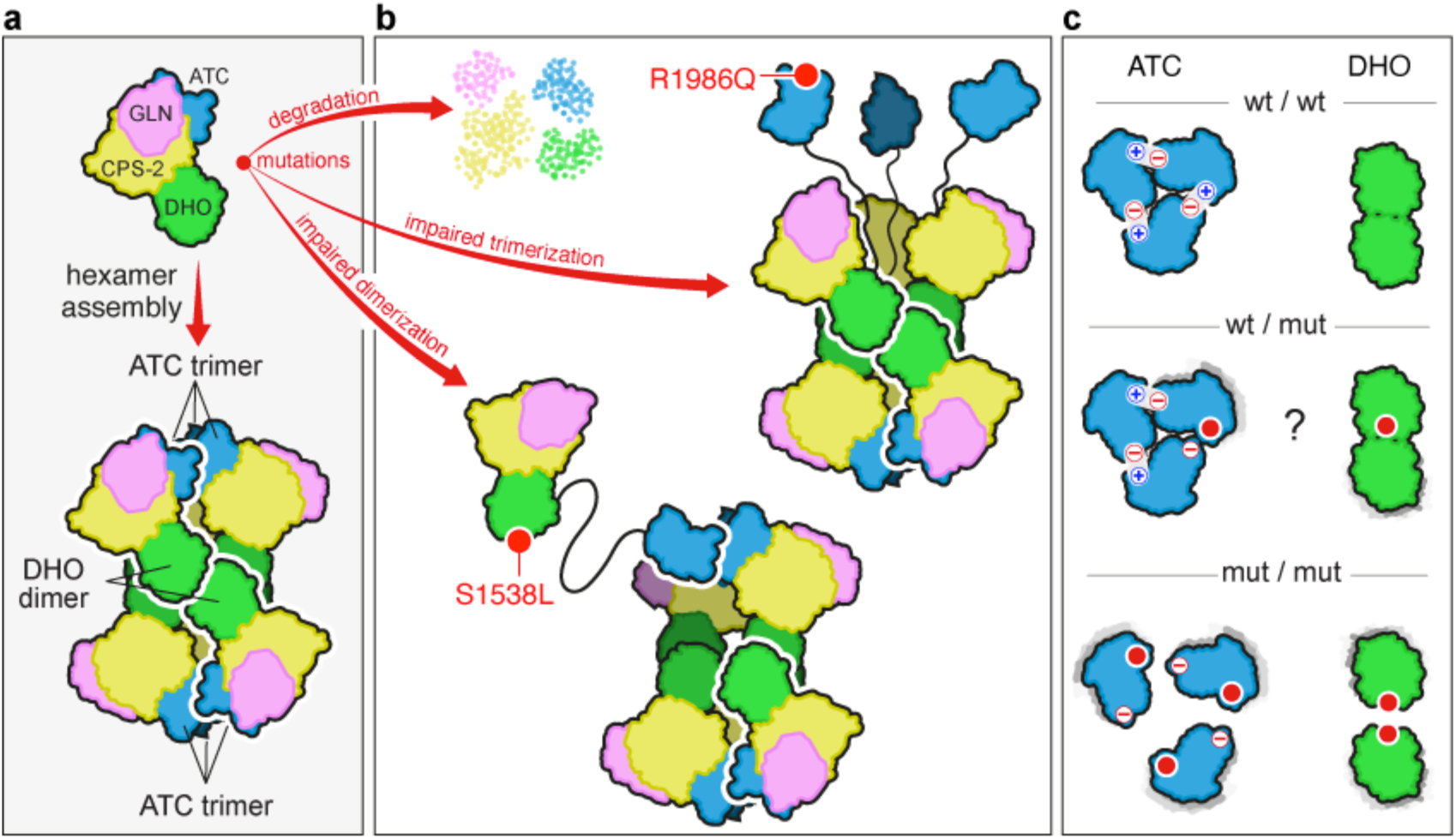
Assembly of CAD pathogenic variants. **a** Proposed model for the assembly of CAD into a hexamer or a “dimer of trimers”. **b** Pathogenic mutations affecting oligomerization (red circles) could lead to protein degradation or “poison” the formation of functional particles. **c** Three scenarios for the oligomerization of WT and mutated ATC and DHO domains. “+” and “-” signs indicate the ion pair between R1986 (+) and D2019 (-).

In summary, the identification and characterization of pathogenic variants has enriched our understanding of CAD, demonstrating that oligomerization into a hexameric complex is essential for its function in the nucleotide biosynthetic pathway. Defects in this assembly process must be carefully considered when assessing the disease-causing potential of new clinical variants. We look forward to experimentally confirming the predicted architecture of this mega-enzyme to shed light on its mechanisms and pathogenic alterations.

## METHODS

### Protein production

The cDNA encoding the human DHO domain (aa 1456–1846) and the ATC domain (aa 1917-2225) in the mutated CAD constructs were PCR amplified using specific primers and inserted into pOPIN-M (Oxford Protein Production Facility) by IVA (in vivo assembly) cloning^47^. WT and mutated DHO were expressed in HEK293 cells and purified as reported^19,20,44^. WT and mutated ATC were expressed in bacteria and purified as previously reported^21^.

### Enzymatic assays

DHO catalyzes a reversible reaction that is pH-dependent^32^. Enzymatic activity was assayed spectrophotometrically following the production of dihydroorotate by absorbance at 230 nm at the favorable acidic pH and at 25 °C^19^. Reactions were carried out in a final volume of 100 μl containing 50 mM sodium phosphate pH 5.7, 150 mM NaCl, 20 μM ZnSO_4_, 0.1 mg·ml^-1^ of bovine serum albumin (BSA; Sigma), and 5 mM carbamoyl aspartate (Ca-Asp; Sigma). Protein concentrations were 0.25 μM for the WT and were increased to 2.5-3 μM for the mutants. The reaction was triggered by adding Ca-Asp to a mix with the enzyme pre-incubated for 5 min at 25 °C.

ATC activity was assayed by a colorimetric method that quantifies the production of Ca-Asp^31,48^. Reactions were carried out in 50 mM Tris-acetate pH 8.3 and 0.1 mg·ml^-1^ BSA in a final volume of 100 µl. Protein was pre-incubated with aspartate for 10 min by floating the tube in a water bath at 25 °C. The reaction was triggered by adding 5 mM carbamoyl phosphate (Sigma) and stopped at different times with 250 µl of color mix solution. The samples were boiled at 95 °C for 15 min and kept in the dark for 30 min before measuring the absorbance at 450 nm. Data analysis was performed with GraphPad Prism.

### Differential scanning fluorimetry (DSF)

Protein stability was measured in triplicates by differential scanning fluorometry^49^ using a CFX Opus 96 Real-Time PCR System (BioRad) in a 96-well reaction plate with 20 μl sample, containing 5 μM protein in GF buffer (20 mM Tris-HCl pH 7, 0.150 M NaCl and 2 mM DTT), 20x SYPRO Orange (Invitrogen) with or without 2 mM 5-fluoroorotate (FOA; Sigma). Fluorescence changes were monitored every 1 °C in a temperature ramp from 20 to 95°C using the extrinsic fluorescence of SYPRO Orange (λ_excitation_ = 465 nm, λ_emission_ = 580 nm). Curves were normalized, and the melting temperature (T_m_) was determined as the midpoint of the unfolding transition. Data were analyzed with GraphPad.

### Crystallization and structure determination

Crystals of the mutated DHO alone or in the presence of 4 mM of Ca-Asp were obtained as reported^19,20,44^. Optimal crystallization conditions consisted of 2-3 mg·ml^-1^ of protein in GF buffer, 2-3 M sodium formate, and 0.1 M HEPES pH 6.5-7.5 as the mother liquor. Crystals were cryoprotected with 20% glycerol and flash-frozen. X-ray diffraction datasets were collected at XALOC (ALBA, Barcelona) and ID23-2 (ESRF, Grenoble). Data processing and scaling were performed automatically with autoPROC^50,51^. Crystallographic phases were obtained by molecular replacement with PHASER^52^ and the WT DHO structure (PDB 4C6C and 4C6J) as the search model. Protein models were traced with COOT^53^ and refined with PHENIX^54^ or Refmac5^55,56^.

### Size-exclusion chromatography coupled to multi-angle light scattering (SEC-MALS) analysis

400 µl of purified protein at 2 mg·ml^-1^ was fractionated on a Superdex 200 10/300 column (Cytiva) equilibrated in GF buffer, using an AKTA purifier (Cytiva). The eluted samples were characterized by in-line measurement of the refractive index and multi-angle light scattering using Optilab T-rEX and DAWN 8+ instruments, respectively (Wyatt Technology). Data were analyzed with ASTRA 6 software^57^ and plotted with GraphPad.

### In silico prediction of pathogenicity

Pathogenicity prediction was performed with AlphaMissense^58^ and FoldX^59^. AlphaMissense scores and pathogenicity predictions were retrieved from https://github.com/google-deepmind/alphamissense. FoldX 5.0 was run locally using default settings, and the models of the crystallographic DHO (PDB ID 4C6C; 1.45 Å resolution) and ATC (PDB 5G1O; 2.1 Å resolution). Before performing calculations, the ‘RepairPDB’ command was used to fix the structures. Individual mutations were introduced using the ‘BuildModel’ command. Calculations were performed in 5 consecutive runs, and ΔΔG values were extracted from the FoldX output files. A value of 1.5 kcal/mol was set as the stability change threshold, as previously suggested in other studies assessing the impact of missense mutations^60,61^.

### Molecular dynamics

Molecular dynamic (MD) simulations were performed for the DHO dimer using GROMACS 2023 software^62^. The DHO WT (PDB ID 4C6I) and S1538L crystal structures were used as input for the solution builder module in CHARMM-GUI^63^ (http://www.charmm-gui.org) and the CHARMM36m^64^ force field was adopted. The system was prepared using the structures without ligands and the flexible loop in the closed conformation, and the non-standard carboxylated lysine residue (K1556) was parameterized with the CHARMM36 force field^65^. Zinc ions were not included in the system due to the complex parameterization of the catalytic center of the protein. The system pH was set to 7, and TIP3 water was added to the simulation in a rectangular box. Potassium and chloride counter-ions in a 0.15 M concentration were added to neutralize the solvated system. The Particle Mesh Ewald (PME)^66^ was used to treat long-range electrostatic interactions, while a 12 Å cutoff scheme was used for van der Waals interactions. Simulations were performed at 310.15 K and 1 bar in an NPT ensemble. Two replicates of 500 ns were run for the WT and mutant dimers, and the trajectories were analyzed individually for each subunit using GROMACS analysis tools. Principal components analysis (PCA) was performed on the backbone atoms to reduce dimensionality and explore protein flexibility along the trajectory. The analysis was performed on single trajectories that concatenated the simulations of the dimer subunits in the two replicas fitted to the reference structure. The covariance matrix and PCA projection for the two first eigenvectors were computed with GROMACS analysis tools. Matplotlib^67^ was used for PCA plot representation, Pymol (https://pymol.org/) and ChimeraX^68^ for visual structural analysis, and GraphPad for other graphical representations.

## ACKNOWLEDGEMENTS

The authors thank the clinicians, Drs. Paula Sanchez Pintos (Instituto de Investigación Sanitaria Santiago de Compostela, La Coruña, Spain) and Joanna Kenny (Children’s Health Ireland at Crumlin, Dublin, Ireland) for providing follow-up on the patients carrying the pathogenic variants reported in this study. This work was supported by grants PID2021-128468NB-I00 and PID2020-120258GB-I00 financed by MCIN/AEI/10.13039/501100011033 to SR-M and RF-L, respectively, and from Fundación Ramón Areces Ciencias de la Vida (XX National Call) to SR-M, and Ramon y Cajal fellowship RYC-2017-23128 to RF-L. LRM was partially supported by the FIB-HCSC intramural young researcher mobility fellowship program 2023, FIB-HCSC, Madrid, Spain. X-ray diffraction experiments at synchrotrons were done through the participation of SR-M in the BAG proposals 2023077633 at ALBA, and MX-2452 at the European Synchrotron Radiation Facility (ESRF) with DOI (10.15151/ESRF-ES-1117952942). The authors thank the ALBA and ESRF synchrotron staff for assistance during data collection.

## AUTHOR CONTRIBUTIONS

FdC and SR-M conceived, planned and performed the experiments. LR-M peformed molecular dynamics simulations with support from LE. MM-M and RF-L contributed to the modeling and interpretation of CAD structure. All authors discussed the results and contributed to the final manuscript.

## DATA AVAILABILITY

Atomic coordinates for S1538L and S1538A mutants are deposited in the Protein Data Bank with PDB IDs: 9FS1 (S1538L bound to Ca-Asp), 9FS2 (S1538A bound to Ca-Asp) and 9FS3 (S1538A in apo form). Previously published structures used in model building and structural comparison are available in the Protein Data Bank with PDB IDs: 5G1N, 5G1O, and 5G1P for ATC and 4C6C, 4C6I, and 4C6M for DHO domains.

## REFERENCES

1. Del Cano-Ochoa F, Moreno-Morcillo M, Ramon-Maiques S (2019) CAD, A Multienzymatic Protein at the Head of de Novo Pyrimidine Biosynthesis. Subcell Biochem 93:505–538.

2. Coleman PF, Suttle DP, Stark GR (1977) Purification from hamster cells of the multifunctional protein that initiates de novo synthesis of pyrimidine nucleotides. The Journal of biological chemistry 252:6379–85.

3. Jones ME (1980) Pyrimidine nucleotide biosynthesis in animals: genes, enzymes, and regulation of UMP biosynthesis. Annual review of biochemistry 49:253–279.

4. Ng BG, Wolfe LA, Ichikawa M, Markello T, He M, Tifft CJ, Gahl WA, Freeze HH (2015) Biallelic mutations in CAD, impair de novo pyrimidine biosynthesis and decrease glycosylation precursors. Human molecular genetics 24:3050–7.

5. Koch J, Mayr JA, Alhaddad B, Rauscher C, Bierau J, Kovacs-Nagy R, Coene KLM, Bader I, Holzhacker M, Prokisch H, et al. (2017) CAD mutations and uridine-responsive epileptic encephalopathy. Brain 140:279–286.

6. Rymen D, Lindhout M, Spanou M, Ashrafzadeh F, Benkel I, Betzler C, Coubes C, Hartmann H, Kaplan JD, Ballhausen D, et al. (2020) Expanding the clinical and genetic spectrum of CAD deficiency: an epileptic encephalopathy treatable with uridine supplementation. Genetics in Medicine [Internet] 22:1589–1597. Available from: https://www.nature.com/articles/s41436-020-0933-z

7. Al-Otaibi A, AlAyed A, Al Madhi A, Saeed L, Ng BG, Freeze HH, Almannai M (2022) Uridine monophosphate (UMP)-responsive developmental and epileptic encephalopathy: A case report of two siblings and a review of literature. Mol Genet Metab Rep 30:100835.

8. Duan L, Ye L, Yin R, Sun Y, Yu W, Zhang Y, Zhong H, Bao X, Tian X (2024) Novel CAD gene mutations in a boy with developmental and epileptic encephalopathy 50 with dramatic response to uridine therapy: a case report and a review of the literature. BMC Pediatrics [Internet] 24:160. Available from: 10.1186/s12887-024-04593-6

9. Frederick A, Sherer K, Nguyen L, Ali S, Garg A, Haas R, Sahagian M, Bui J (2021) Triacetyluridine treats epileptic encephalopathy from CAD mutations: a case report and review. Annals of Clinical and Translational Neurology [Internet] 8:284–287. Available from: https://onlinelibrary.wiley.com/doi/abs/10.1002/acn3.51257

10. Kamate M, Patil S (2020) CAD Deficiency—Another Treatable Early Infantile Epileptic Encephalopathy. Pediatric Neurology [Internet] 110:97–98. Available from: https://www.pedneur.com/article/S0887-8994(20)30151-X/abstract

11. McGraw CM, Mahida S, Jayakar P, Koh HY, Taylor A, Resnick T, Rodan L, Schwartz MA, Ejaz A, Sankaran VG, et al. (2021) Uridine-responsive epileptic encephalopathy due to inherited variants in CAD: A Tale of Two Siblings. Annals of Clinical and Translational Neurology [Internet] 8:716–722. Available from: https://onlinelibrary.wiley.com/doi/abs/10.1002/acn3.51272

12. Peng X, Xia L, Zhang H, Zhang J, Yu S, Wang S, Xu Y, Yao B, Ye J (2022) A Treatable Genetic Disease Caused by CAD Mutation. Front. Pediatr. [Internet] 10. Available from: https://www.frontiersin.org/articles/10.3389/fped.2022.771374

13. Steinberg-Shemer O, Yacobovich J, Noy-Lotan S, Dgany O, Krasnov T, Barg A, Landau YE, Kneller K, Somech R, Gilad O, et al. (2023) Biallelic hypomorphic variants in CAD cause uridine-responsive macrocytic anaemia with elevated haemoglobin-A2. Br J Haematol.

14. Yarahmadi SG, Morovvati S (2022) CAD gene and early infantile epileptic encephalopathy-50; three Iranian deceased patients and a novel mutation: case report. BMC Pediatrics [Internet] 22:125. Available from: 10.1186/s12887-022-03195-4

15. Zhou L, Xu H, Wang T, Wu Y (2020) A patient with CAD deficiency responsive to uridine and lietrature review. Front. Neurol. 11:5.

16. Zhou L, Xu H, Wang T, Wu Y (2020) A patient with CAD deficiency responsive to uridine and lietrature review. Front. Neurol. 11:5.

17. Del Caño-Ochoa F, Ramón-Maiques S (2021) Deciphering CAD: Structure and function of a mega-enzymatic pyrimidine factory in health and disease. Protein Sci 30:1995–2008.

18. Lee L, Kelly RE, Pastra-Landis SC, Evans DR (1985) Oligomeric structure of the multifunctional protein CAD that initiates pyrimidine biosynthesis in mammalian cells. Proc Natl Acad Sci U S A 82:6802–6.

19. Grande-Garcia A, Lallous N, Diaz-Tejada C, Ramon-Maiques S (2014) Structure, functional characterization, and evolution of the dihydroorotase domain of human CAD. Structure 22:185–98.

20. Lallous N, Grande-Garcia A, Molina R, Ramon-Maiques S (2012) Expression, purification, crystallization and preliminary X-ray diffraction analysis of the dihydroorotase domain of human CAD. Acta crystallographica. Section F, Structural biology and crystallization communications 68:1341–5.

21. Ruiz-Ramos A, Velazquez-Campoy A, Grande-Garcia A, Moreno-Morcillo M, Ramon-Maiques S (2016) Structure and Functional Characterization of Human Aspartate Transcarbamoylase, the Target of the Anti-tumoral Drug PALA. Structure 24:1081–94.

22. Ruiz-Ramos A, Lallous N, Grande-Garcia A, Ramon-Maiques S (2013) Expression, purification, crystallization and preliminary X-ray diffraction analysis of the aspartate transcarbamoylase domain of human CAD. Acta crystallographica. Section F, Structural biology and crystallization communications 69:1425–30.

23. Carrey EA The shape of CAD. In: Davidson JN, editor. Paths to pyrimidines - an international newsletter. Vol. 3. University of Kentuky; 1995. pp. 68–72.

24. Moreno-Morcillo M, Grande-Garcia A, Ruiz-Ramos A, Del Cano-Ochoa F, Boskovic J, Ramon-Maiques S (2017) Structural Insight into the Core of CAD, the Multifunctional Protein Leading De Novo Pyrimidine Biosynthesis. Structure 25:912–923 e5.

25. Del Caño-Ochoa F, Ng BG, Abedalthagafi M, Almannai M, Cohn RD, Costain G, Elpeleg O, Houlden H, Karimiani EG, Liu P, et al. (2020) Cell-based analysis of CAD variants identifies individuals likely to benefit from uridine therapy. Genet Med 22:1598–1605.

26. Del Caño-Ochoa F, Ng BG, Rubio-del-Campo A, Mahajan S, Wilson MP, Vilar M, Rymen D, Sánchez-Pintos P, Kenny J, Ley Martos M, et al. (2023) Beyond genetics: Deciphering the impact of missense variants in CAD deficiency. J of Inher Metab Disea:jimd.12667.

27. Collins KD, Stark GR (1971) Aspartate transcarbamylase interaction with the transition state analogue N- (phosphonacetyl)-L-aspartate. Journal of Biological Chemistry 246:6599–6605.

28. Jumper J, Evans R, Pritzel A, Green T, Figurnov M, Ronneberger O, Tunyasuvunakool K, Bates R, Žídek A, Potapenko A, et al. (2021) Highly accurate protein structure prediction with AlphaFold. Nature [Internet] 596:583–589. Available from: https://www.nature.com/articles/s41586-021-03819-2

29. Thoden JB, Holden HM, Wesenberg G, Raushel FM, Rayment I (1997) Structure of carbamoyl phosphate synthetase: a journey of 96 A from substrate to product. Biochemistry 36:6305–16.

30. de Cima S, Polo LM, Diez-Fernandez C, Martinez AI, Cervera J, Fita I, Rubio V (2015) Structure of human carbamoyl phosphate synthetase: deciphering the on/off switch of human ureagenesis. Sci Rep 5:16950.

31. Ruiz-Ramos A, Velazquez-Campoy A, Grande-Garcia A, Moreno-Morcillo M, Ramon-Maiques S (2016) Structure and Functional Characterization of Human Aspartate Transcarbamoylase, the Target of the Anti-tumoral Drug PALA. Structure 24:1081–94.

32. Christopherson RI, Jones ME (1980) The overall synthesis of L-5, 6-dihydroorotate by multienzymatic protein pyr1-3 from hamster cells. Kinetic studies, substrate channeling, and the effects of inhibitors. J. Biol. Chem. 255:11381–11395.

33. Otsuki T, Mori M, Tatibana M (1982) Studies on channeling of carbamoyl-phosphate in the multienzyme complex that initiates pyrimidine biosynthesis in rat ascites hepatoma cells. J Biochem 92:1431–7.

34. Mally MI, Grayson DR, Evans DR (1980) Catalytic synergy in the multifunctional protein that initiates pyrimidine biosynthesis in Syrian hamster cells. The Journal of biological chemistry 255:11372–80.

35. Carrey EA, Campbell DG, Hardie DG (1985) Phosphorylation and activation of hamster carbamyl phosphate synthetase II by cAMP-dependent protein kinase. A novel mechanism for regulation of pyrimidine nucleotide biosynthesis. The EMBO journal 4:3735.

36. Ben-Sahra I, Howell JJ, Asara JM, Manning BD (2013) Stimulation of de novo pyrimidine synthesis by growth signaling through mTOR and S6K1. Science 339:1323–8.

37. Robitaille AM, Christen S, Shimobayashi M, Cornu M, Fava LL, Moes S, Prescianotto-Baschong C, Sauer U, Jenoe P, Hall MN (2013) Quantitative phosphoproteomics reveal mTORC1 activates de novo pyrimidine synthesis. Science 339:1320–3.

38. Graves LM, Guy HI, Kozlowski P, Huang M, Lazarowski E, Pope RM, Collins MA, Dahlstrand EN, Earp HS, Evans DR (2000) Regulation of carbamoyl phosphate synthetase by MAP kinase. Nature 403:328–32.

39. Yang C, Zhao Y, Wang L, Guo Z, Ma L, Yang R, Wu Y, Li X, Niu J, Chu Q, et al. (2023) De novo pyrimidine biosynthetic complexes support cancer cell proliferation and ferroptosis defence. Nat Cell Biol 25:836–847.

40. Qiu Y, Davidson JN (2000) Substitutions in the aspartate transcarbamoylase domain of hamster CAD disrupt oligomeric structure. Proc Natl Acad Sci U S A 97:97–102.

41. Qiu Y, Davidson JN (1998) Aspartate-90 and arginine-269 of hamster aspartate transcarbamylase affect the oligomeric state of a chimaeric protein with an Escherichia coli maltose-binding domain. The Biochemical journal 329 (Pt 2):243–7.

42. Davidson JN, Rumsby PC, Tamaren J (1981) Organization of a multifunctional protein in pyrimidine biosynthesis. Analyses of active, tryptic fragments. The Journal of biological chemistry 256:5220–5.

43. Souciet JL, Nagy M, Le Gouar M, Lacroute F, Potier S (1989) Organization of the yeast URA2 gene: identification of a defective dihydroorotase-like domain in the multifunctional carbamoylphosphate synthetase-aspartate transcarbamylase complex. Gene 79:59–70.

44. Del Cano-Ochoa F, Grande-Garcia A, Reverte-Lopez M, D’Abramo M, Ramon-Maiques S (2018) Characterization of the catalytic flexible loop in the dihydroorotase domain of the human multi-enzymatic protein CAD. J Biol Chem 293:18903–18913.

45. Lee M, Maher MJ, Christopherson RI, Guss JM (2007) Kinetic and structural analysis of mutant Escherichia coli dihydroorotases: a flexible loop stabilizes the transition state. Biochemistry 46:10538–50.

46. Lee M, Chan CW, Mitchell Guss J, Christopherson RI, Maher MJ (2005) Dihydroorotase from Escherichia coli: loop movement and cooperativity between subunits. J Mol Biol 348:523–33.

47. García-Nafría J, Watson JF, Greger IH (2016) IVA cloning: A single-tube universal cloning system exploiting bacterial In Vivo Assembly. Sci Rep 6:27459.

48. Prescott LM, Jones ME (1969) Modified methods for the determination of carbamyl aspartate. Anal Biochem 32:408–19.

49. Niesen FH, Berglund H, Vedadi M (2007) The use of differential scanning fluorimetry to detect ligand interactions that promote protein stability. Nat Protoc 2:2212–2221.

50. Monaco S, Gordon E, Bowler MW, Delagenière S, Guijarro M, Spruce D, Svensson O, McSweeney SM, McCarthy AA, Leonard G, et al. (2013) Automatic processing of macromolecular crystallography X-ray diffraction data at the ESRF. J Appl Crystallogr 46:804–810.

51. Vonrhein C, Flensburg C, Keller P, Sharff A, Smart O, Paciorek W, Womack T, Bricogne G (2011) Data processing and analysis with the autoPROC toolbox. Acta Crystallogr D Biol Crystallogr 67:293–302.

52. McCoy AJ, Grosse-Kunstleve RW, Adams PD, Winn MD, Storoni LC, Read RJ (2007) Phaser crystallographic software. J Appl Crystallogr 40:658–674.

53. Emsley P, Lohkamp B, Scott WG, Cowtan K (2010) Features and development of Coot. Acta Crystallogr D Biol Crystallogr 66:486–501.

54. Adams PD, Afonine PV, Bunkoczi G, Chen VB, Davis IW, Echols N, Headd JJ, Hung LW, Kapral GJ, Grosse-Kunstleve RW, et al. (2010) PHENIX: a comprehensive Python-based system for macromolecular structure solution. Acta Crystallogr D Biol Crystallogr 66:213–21.

55. Murshudov GN, Skubak P, Lebedev AA, Pannu NS, Steiner RA, Nicholls RA, Winn MD, Long F, Vagin AA (2011) REFMAC5 for the refinement of macromolecular crystal structures. Acta Crystallogr D Biol Crystallogr 67:355– 67.

56. Winn MD, Ballard CC, Cowtan KD, Dodson EJ, Emsley P, Evans PR, Keegan RM, Krissinel EB, Leslie AG, McCoy A, et al. (2011) Overview of the CCP4 suite and current developments. Acta Crystallogr D Biol Crystallogr 67:235–42.

57. Wyatt PJ (1993) Light scattering and the absolute characterization of macromolecules. Analytica Chimica Acta 272:1–40.

58. Cheng J, Novati G, Pan J, Bycroft C, Žemgulytė A, Applebaum T, Pritzel A, Wong LH, Zielinski M, Sargeant T, et al. (2023) Accurate proteome-wide missense variant effect prediction with AlphaMissense. Science [Internet] 381:eadg7492. Available from: https://www.science.org/doi/10.1126/science.adg7492

59. Schymkowitz J, Borg J, Stricher F, Nys R, Rousseau F, Serrano L (2005) The FoldX web server: an online force field. Nucleic Acids Research [Internet] 33:W382–W388. Available from: 10.1093/nar/gki387

60. Bromberg Y, Rost B (2009) Correlating protein function and stability through the analysis of single amino acid substitutions. BMC Bioinformatics [Internet] 10:S8. Available from: 10.1186/1471-2105-10-S8-S8

61. Seifi M, Walter MA (2018) Accurate prediction of functional, structural, and stability changes in PITX2 mutations using in silico bioinformatics algorithms. PLOS ONE [Internet] 13:e0195971. Available from: https://journals.plos.org/plosone/article?id=10.1371/journal.pone.0195971

62. Abraham MJ, Murtola T, Schulz R, Páll S, Smith JC, Hess B, Lindahl E (2015) GROMACS: High performance molecular simulations through multi-level parallelism from laptops to supercomputers. SoftwareX [Internet] 1– 2:19–25. Available from: https://www.sciencedirect.com/science/article/pii/S2352711015000059

63. Jo S, Kim T, Iyer VG, Im W (2008) CHARMM-GUI: A web-based graphical user interface for CHARMM. Journal of Computational Chemistry [Internet] 29:1859–1865. Available from: https://onlinelibrary.wiley.com/doi/abs/10.1002/jcc.20945

64. Huang J, Rauscher S, Nawrocki G, Ran T, Feig M, de Groot BL, Grubmüller H, MacKerell AD (2017) CHARMM36m: an improved force field for folded and intrinsically disordered proteins. Nat Methods [Internet] 14:71–73. Available from: https://www.nature.com/articles/nmeth.4067

65. Croitoru A, Park S-J, Kumar A, Lee J, Im W, MacKerell AD Jr, Aleksandrov A (2021) Additive CHARMM36 Force Field for Nonstandard Amino Acids. J. Chem. Theory Comput. [Internet] 17:3554–3570. Available from: 10.1021/acs.jctc.1c00254

66. Darden T, Tork D, Pedersen L (1997) Particle mesh Ewald: An N-log(N) method for Ewald sums in large systems. J. Comput. Chem. 18:1463–1472.

67. Hunter J (2007) Matplotlib: A 2D Graphics Environment. Computing in Science & Engineering 9:90–95.

68. Pettersen EF, Goddard TD, Huang CC, Couch GS, Greenblatt DM, Meng EC, Ferrin TE (2004) UCSF Chimera--a visualization system for exploratory research and analysis. J Comput Chem 25:1605–12.

