## Supplementary Figures for "Disruption of CAD oligomerization by pathogenic variants"

---

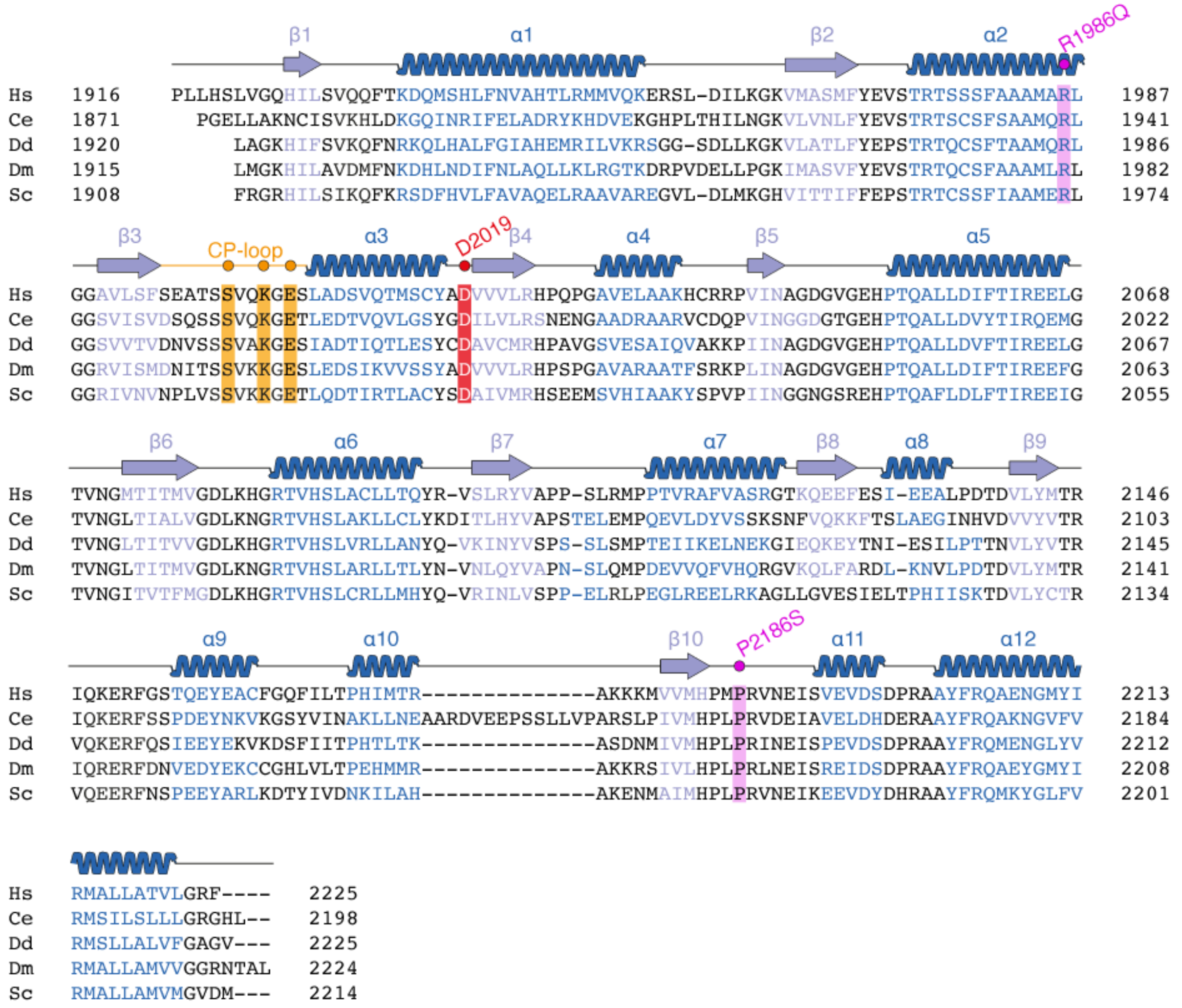


**Supplementary Figure 1. Sequence alignment of CAD's ATC domain.** Alignment of ATC domain sequences from human (Hs, UniProt P27708), *Caenorhabditis elegans* (Ce, UniProt Q18990), *Dictyostelium discoideum* (Dd, P20054), *Drosophila melanogaster* (Dm, P05990) and *Saccharomyces cerevisiae* (Sc, P07259). Secondary structure elements of human ATC are indicated as arrows (β-strands) and spirals (α-helices) above the sequence. The CP-loop is highlighted in orange. Positions affected by pathogenic mutations (R1986Q and P2186S) are highlighted in magenta. The conserved aspartate forming the ionic pair (D2019) and key residues in the CP-loop are colored red and orange, respectively.

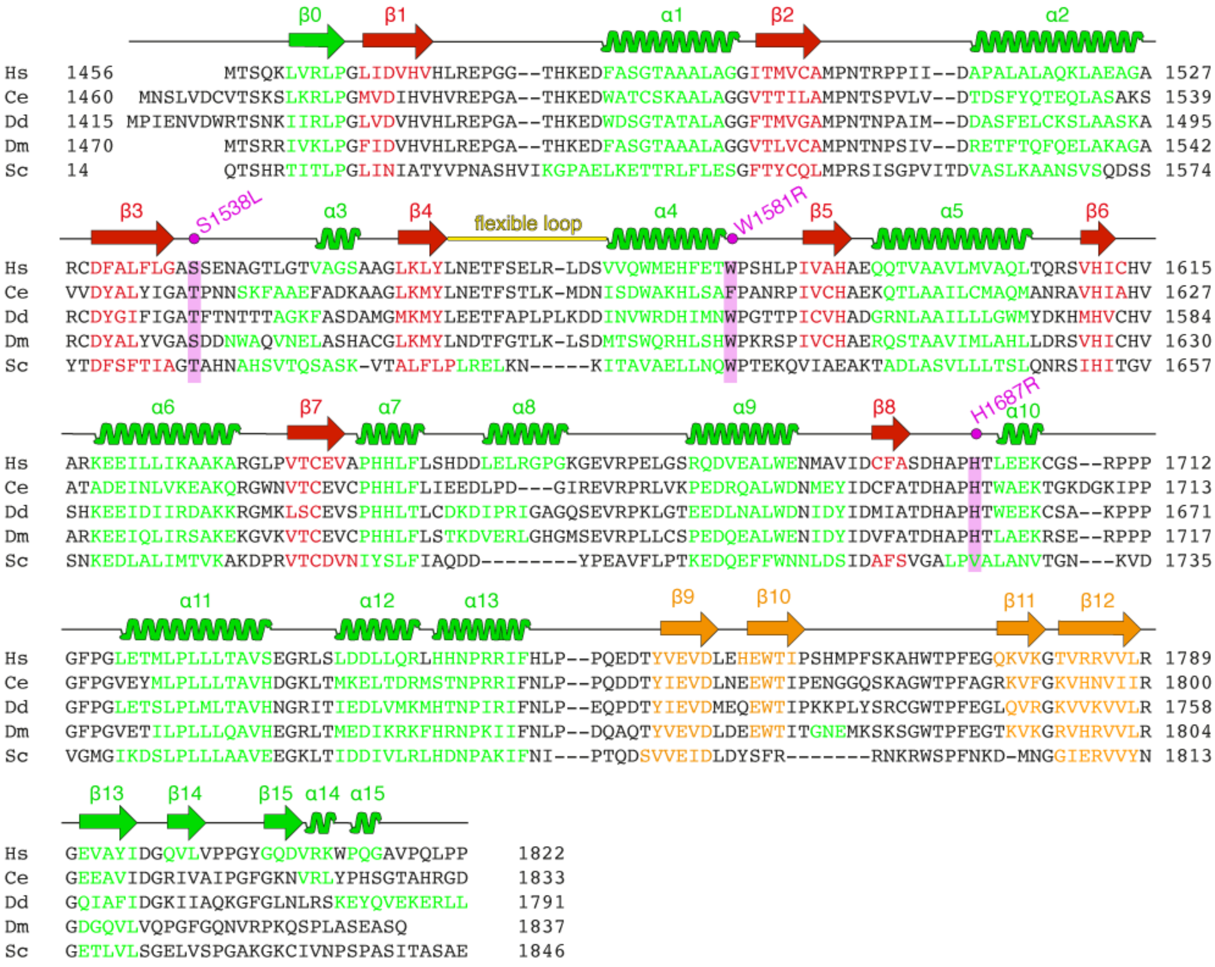


**Supplementary Figure 2. Sequence alignment of CAD's DHO domain.** Alignment of DHO domain sequences from human (Hs, UniProt P27708), *Caenorhabditis elegans* (Ce, UniProt Q18990), *Dictyostelium discoideum* (Dd, P20054), *Drosophila melanogaster* (Dm, P05990) and *Saccharomyces cerevisiae* (Sc, P07259). Secondary structure elements of human DHO are indicated as arrows (β-strands) and spirals (α-helices) above the sequence. The catalytic flexible loop is highlighted in yellow. Positions affected by pathogenic mutations (S1538L, W1581R and H1687R) are highlighted in magenta.

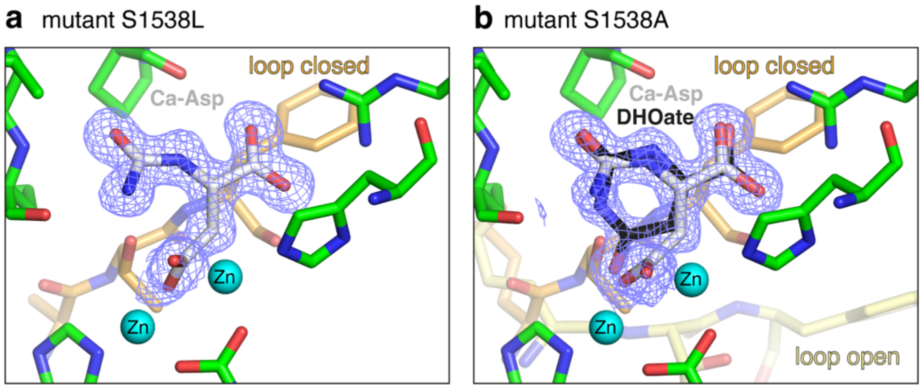


**Supplementary Figure 3. Ligand binding in the crystal structures of DHO mutants S1538L and S1538A**. Detailed view of the active sites of DHO mutants S1538L (**a**) and S1538A (**b**). The $2F_{obs}$-$F_{calc}$ electron density map for the ligand is shown as a blue mesh contoured at 1.0 σ. Ca-Asp and dihydroorotate (DHOate) are depicted with carbon atoms in grey or black, respectively. $Zn^{2+}$ ions are represented as cyan spheres. The flexible loop is orange and yellow for the open and closed states.

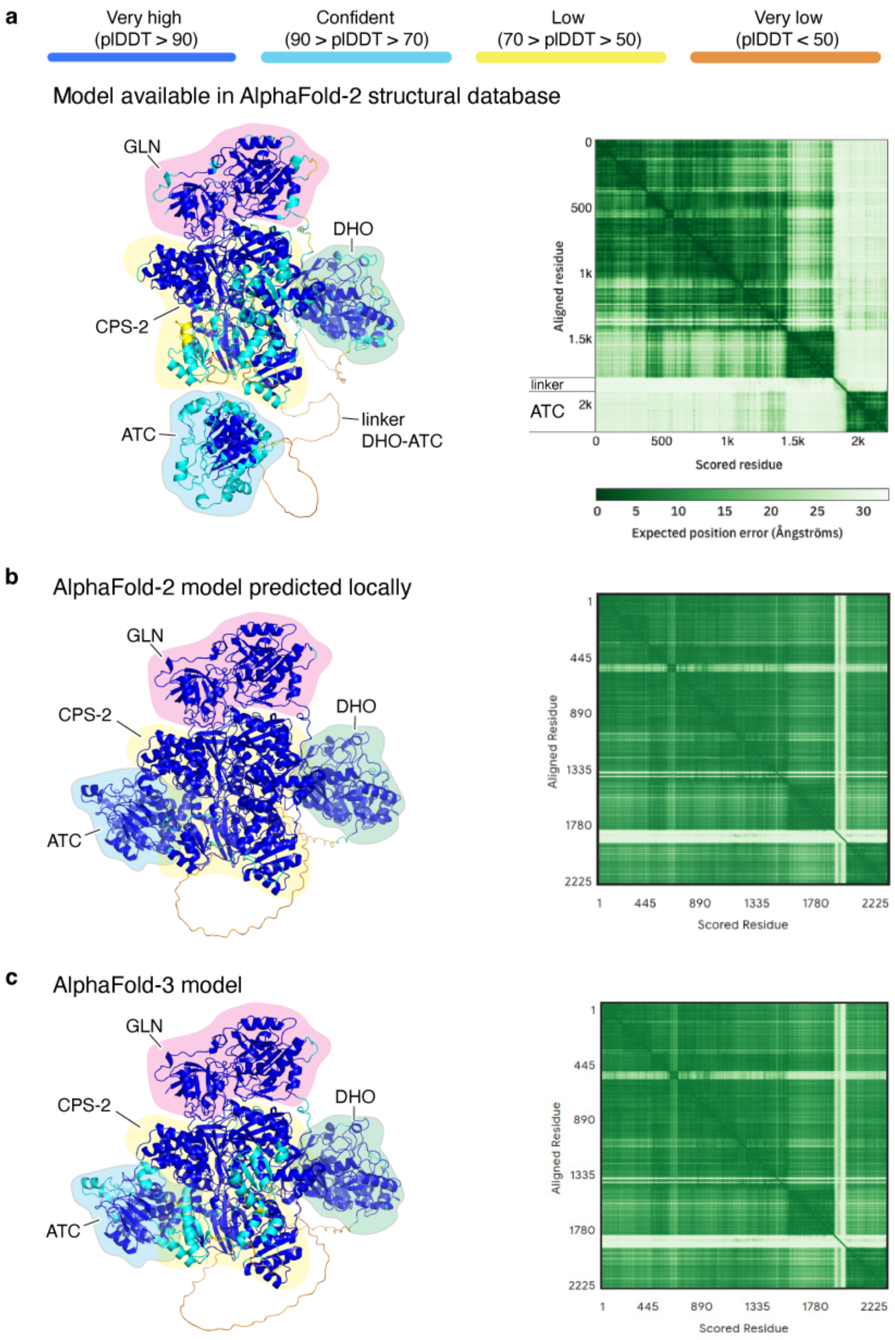


**Supplementary Figure 4. Structural models of CAD subunit predicted by AlphaFold.** Cartoon representation of AlphaFold models available in the database (**a**), obtained by running AlphaFold-2 locally (**b**), or from AlphaFold-3 (**c**). The structures are colored according to the per-atom confidence accuracy value (pLDDT). The predicted aligned error (PAE) plots are shown for each model.

**Supplementary Table 1.** *In silico* pathogenicity predictions for DHO and ATC missense mutations.

| | | AlphaMissense | | FoldX | |
|---|---|---|---|---|---|
| | Mutation | Score[a] | Prediction | ΔΔG (kcal/mol)[b] | Prediction |
| DHO | S1538L | 0,325 | benign | 3,486 | pathogenic |
| | W1581R | 0,995 | pathogenic | 7,382 | pathogenic |
| | H1687R | 0,948 | pathogenic | 7,024 | pathogenic |
| ATC | R1986Q | 0,879 | pathogenic | 3,008 | pathogenic |
| | P2186S | 0,995 | pathogenic | 3,108 | pathogenic |

[a]AlphaMissense scores retrieved from https://github.com/google-deepmind/alphamissense. Higher values indicate increased pathogenicity prediction.

[b]FoldX ΔΔG is the difference between $\Delta G^{Mutant}$ and $\Delta G^{WT}$ and is computed as the mean score per subunit in each domain. A mutation is considered destabilizing if it results in higher energy than the WT. A value of 1.5 kcal/mol was set as the stability change threshold.
